# Discovery of non-canonical proteins through modification-aware proteogenomics

**DOI:** 10.64898/2026.08.17.745157

**Authors:** Valeriia Vasylieva, Enrico Massignani, Tine Claeys, Francis Bourassa, Sebastien Leblanc, Ihor Arefiev, Lennart Martens, Marie A. Brunet

**Author notes:** co-first (Valeriia Vasylieva, Enrico Massignani).

## Abstract

**Short:** The SwissProt database contains a stable 20,418 human protein-coding genes and 42,541 human protein sequences. Ribo-Seq suggests about 7,000 additional, non-canonical Open Reading Frames (ORFs) are present in humans, though only a few of them are confirmed by Mass Spectrometry (MS). Detecting these proteins requires extensive database searches, increasing computational load and inflating False Discovery Rates (FDR). Using the ionbot search engine with the OpenProt database allows for reliable detection of non-canonical proteins while controlling FDR. Ionbot surpasses the Trans-Proteomics Pipeline (TPP) in reproducibility, identifying more peptides and proteins supported by multiple spectra. In addition, open modification searches yield better PSMs compared to closed searches. This work highlights the importance of employing cutting-edge search engines in non-canonical protein research, as well as the value of open modification search in correcting errors in non-canonical protein detection.

**Long:** *Background:* The SwissProt database reports a quite stable 20,418 human protein-coding genes and 42,541 human protein sequences, figures that have remained stable. New techniques like Ribo-Seq indicate that approximately 7,000 additional, non-canonical Open Reading Frames (ORFs) are translated in humans, few of which have been confirmed by Mass Spectrometry (MS). Detecting these non-canonical proteins requires comprehensive database searches, which increase computational load and False Discovery Rate (FDR). Here, we use the open search engine ionbot in combination with the OpenProt proteogenomics database to reproducibly detect non-canonical proteins while maintaining a well-controlled FDR.

*Results:* Compared to the current gold standard, the Trans-Proteomics Pipeline (TPP), ionbot shows higher reproducibility, with a higher number of peptides and proteins supported by multiple spectra, and across multiple samples. We observe that PSMs from the open modification search against OpenProt have higher fragment ion intensity correlation compared to PSMs obtained from the closed search, or by only searching canonical proteins.

*Conclusions:* In this work, we show the potential for open modification searching to correct potential mistakes in non-canonical proteins detection by preventing modified canonical peptides or variants from being incorrectly identified as non-canonical peptides. We also highlight the importance of assessing the FDR of non-canonical identifications separately from canonical ones, as global FDR calculations are biased by the scarcity of non-canonical identifications in each dataset.

## Background

Liquid chromatography-tandem mass spectrometry (LC-MS/MS) is the gold standard for high-throughput proteomics, enabling the identification and quantification of thousands of proteins in a single sample, providing a comprehensive view of the proteome and its complexity [1]. The vast amounts of spectral data generated by LC-MS/MS require computational processing using peptide search engines. The most common approach relies on database search engines, which match experimental MS/MS spectra against theoretical spectra generated through *in silico* digestion of a reference proteome [2].

A fundamental limitation of database searching is its dependence on search space size. As the database grows, the probability of incorrect peptide-spectrum matches (PSMs) with artificially high scores increases [3]. This can inflate the false discovery rate (FDR) and lower the overall identification rates. Search space size is influenced by several factors, the most significant being the protein database itself. While limiting searches to tryptic peptides from annotated proteins can improve efficiency, it is impossible for applications such as peptidomics, which aims to identify non-tryptic peptides, or for proteogenomics.

Proteogenomics involves searching a database of proteins predicted from genomic data. Theoretical proteins can be derived from open reading frames of custom genomes, narrowed down to captured mRNAs or translation events recorded by Ribo-seq. The classical proteomics approach, in contrast, relies on reference proteomes, but these do not account for amino acid variants, alternative translation start sites, or non-canonical proteins. These latter proteins, so called due to their absence from conventional proteomics databases like UniProtKB, can be proteins derived from small open reading frames, dual-coding genes, pseudogenes, or unrecorded isoforms. Lower expression levels and shorter sequences of these non-canonical proteins reduce their likelihood of producing detectable peptides [4,5]. Moreover, non-canonical proteins also produce fewer proteotypic peptides than canonical proteins, making protein inference more challenging [5].

According to OpenProt, a proteogenomics resource aimed at cataloguing non-canonical proteins, 13.8% of identifiable tryptic peptides are shared between canonical and non-canonical proteins, and 6.3% of non-canonical proteins lack proteotypic peptides altogether under trypsin digestion. Reproducible detection remains a concern for non-canonical proteins, which are often debated to be spurious or unstable gene products. Nevertheless, numerous non-canonical proteins have been validated *in vivo* and have been demonstrated to be involved in various biological processes [6].

Beyond non-canonical proteins, the search space can be further expanded by the inclusion of post-translational modifications (PTMs). Indeed, to accurately account for modifications, search engines must generate theoretical spectra for both unmodified and all possible modified forms of each peptide. However, if a given PTM is not included in the search, but is present on proteins in the sample, peptides modified by that PTM can either be misidentified as different, unmodified peptides, or remain unidentified altogether. Indeed, a substantial portion of unidentified spectra comes from modified spectra [7-9].

Here, we therefore hypothesize that open modification search (OMS) can rescue non-canonical peptides with modified residues. Non-canonical peptides with sufficiently different sequences from other canonical peptides, or including unexpected modifications, would be unidentified in closed search pipelines. At the same time, non-canonical protein identifications should be carefully examined to avoid false claims of novel proteins. On top of the quality control steps outlined in Human Proteome Project guidelines, the issue of isobaric precursors is highlighted in open search [10]. Concretely, 82,226 proteotypic, non-canonical protein-derived peptides differ from common peptides by only one amino acid in OpenProt. This small difference in sequence can be mistaken for the presence of unaccounted modifications that are isobaric to amino acid substitutions: for example, Alanine and Valine, or Aspartic Acid and Glutamic Acid, both differ by exactly one methyl group (+14.02 Da). Thus, a protein methylation event on one site can be easily misinterpreted as an amino acid substitution on a nearby site (and *vice versa*). In such cases, it is easy for the search engine to confuse the best peptide sequence match [9], which can affect the trustworthiness of non-canonical protein detections.

However, when proteogenomics and OMS are combined in one study, the search space increases dramatically, which can in turn cause potentially drastic FDR. While each database search engine differs in its strategy to control FDR, a common strategy for improving identification accuracy is the re-scoring of PSMs using tools such as PeptideProphet [11], Percolator [12], or MS2Rescore [13]. However, these approaches typically operate only on the highest-ranked PSMs from the initial search, meaning an erroneous top-ranked PSM cannot be substituted with the correct identification. Another issue with proteogenomics is that the target-decoy approach, which is used to control FDR in proteomics analysis [14–16], assumes that each experimental spectrum has an equal probability to match a false peptide (decoy) or a true one (target). However, this assumption does not always match reality [17], as was observed for peptides grouped by physical properties (e.g. charge) [18], modification state (modified vs unmodified) [19], protein type (canonical vs non-canonical) [5,20], or by proteins sharing the same peptides [21]. Thus, it is recommended that q-values be recalibrated according to the proportion of decoys in each group. It can be done through the application of independent q-value thresholds to each group [20] or with simultaneous control on global FDR with the Group-walk algorithm [22].

To address these challenges, we employed ionbot, a semi-supervised machine learning-based search engine [23]. A key component in ionbot is the concept of a candidate match set, which extends beyond the first-ranked matches and is derived from an extensive open search space using predicted sequence tags and a set of biased PSM scoring functions. Ionbot integrates advanced predictive models for retention time and fragmentation patterns into its scoring function. This combination of sequence tags and sophisticated scoring enhances its peptide matching capabilities. Ionbot showed a higher identification rate compared to MSFragger [24] and open-pFind [25] using a small database (e.g., reference proteome). The correctly estimated FDR allowed ionbot to identify a substantial number of co-eluting matches. It is especially relevant for non-canonical peptides with charge-to-mass ratios similar to canonical peptides. The PSM scoring function used by ionbot is derived from the candidate match set using semi-supervised machine learning, ensuring that the PSM scores are reproducibly tailored to the experimental data. This results in a search engine that surpasses traditional search engines and facilitates reliable open modification searches that exceed the performance of current open modification engines.

In this study, we compare the classical pipelines for closed (Trans Proteomic Pipeline) and open search (MSFragger in the FragPipe pipeline) with ionbot in the context of increasing database size and determine the effectiveness of ionbot FDR control relative to the search space size [26,27]. Based on this endeavor, we apply best practices in the quality control of non-canonical proteins identification.

## Results

### Ionbot ensures more robust FDP control than TPP and FragPipe in proteogenomics searches

Ionbot controls FDR by target-decoy competition, with concatenated-decoy search. However, FDR only gives an estimate of the real fraction of false positives in the search results (known as False Discovery Proportion, or FDP). To assess the FDP, we performed entrapment analyses, in which peptide sequences that should not be present in the biological sample are added to the database [28,29]. Unlike decoy sequences, these new sequences will be treated by the search engines as “targets” for the sake of FDP computation. Here, we adopt FDRBench to compute the upper bounds of the FDP on peptide level at 1% FDR [30].

We examined whether ionbot FDP control varies depending on the search mode and database. The results have been compared to Trans Proteomic Pipeline (TPP) and FragPipe performance. We re-analyzed three public PRIDE datasets, which include MS data derived from breast cancer cell lines (SK-BR-3, MCF7, BT474), a cervical carcinoma cell line (HeLa), and prostate cancer tissue and corresponding lymph node metastases [31-33]. Despite variability among datasets, assessing individual biological contexts is beyond this study. Thus, our conclusions are derived from the combined results of all three datasets. MS raw data was searched against three databases: (i) UniprotKB/Swiss-Prot (reviewed reference proteins); (ii) UniProtKB/TrEMBL (reference proteins, their isoforms and unreviewed sequences); and (iii) OpenProt (includes TrEMBL and non-canonical proteins) [35]. SwissProt, TrEMBL, and OpenProt have size ratios 1 : 5 : 33.5 and are hierarchical, with each subsequent database substantially overlapping the previous one, thus increasing peptide degeneracy. We will use “canonical peptides”, “non-canonical peptides”, and “common peptides” to refer to peptides that could be mapped exclusively to canonical proteins, exclusively to non-canonical proteins, or to both canonical and non-canonical proteins, respectively. The term “peptides of canonical proteins” refers to “canonical” and “common” simultaneously, as non-canonical proteins can only be identified with non-canonical peptides.

The estimated FDP level at 1% FDR was consistent for all databases, showing that ionbot FDP, as well as classic pipelines’ FDP, is not affected by database size, and thus remains well-controlled (Figure 1A). The ionbot estimated FDP reaches a maximum of 2.27% for closed search and 1.9% for open search. Ionbot in closed search (for which it was not designed) mode yields a slightly higher FDP than TPP (maximum = 1.65%). In its intended open search mode, however, ionbot offers much better FDR control, maintaining its FDP 2.7 times lower than FragPipe on average. Ionbot is successful in FDR control for the peptides of canonical proteins (FDP < 1.1%) but fails for the peptides unique to non-canonical proteins (FDP<65.9%) (Figure 1B). The non-canonical peptides FDP is even higher for TPP (maximum = 78.41%) and FragPipe (maximum = 70.02%). Overall, this hints that more diverse or sparse peptide populations (such as non-canonical peptides, which are less likely to be detected due to LC-MS/MS sensitivity limits) present a different distribution of target and decoy PSM scores than the much more abundant canonical ones.

**Figure 1.**
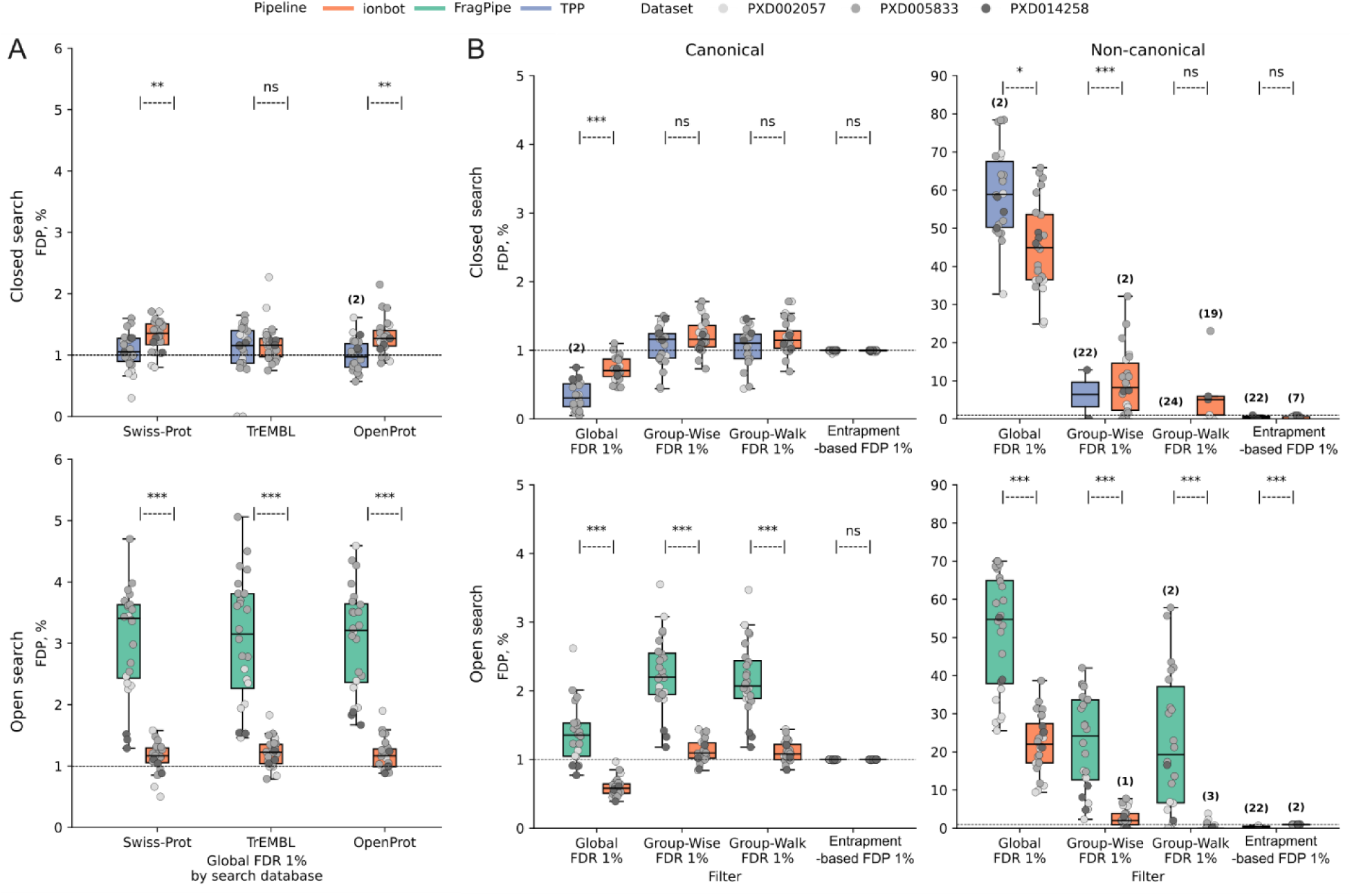
FDR control by classic and ionbot pipelines on peptide level and outcome of filtering strategies. **A.** FDP at Global FDR 1% estimated by FDRBench for each search database. Each dot represents one of the samples from three datasets. Only samples with identifications after filtering are shown. Number of samples without identifications after filtering is indicated above boxplots when applicable. Significance of the difference in estimated FDP between classic pipelines and ionbot is calculated using the Wilcoxon test across all samples (ns – not significant, * p < 0.05, ** p < 0.01, *** p < 0.001). **B.** Estimated FDP per peptide type at FDR 1% for global and group FDR filtering strategies and at FDP 1% for entrapment-based filtering. This analysis was only conducted on results of searches against OpenProt. The estimated FDP of non-canonical peptides identified by ionbot is ∼23% higher in closed search than in open search for global filtering (Wilcoxon p<0.001) and remains higher than 1% even when grouped FDR control is applied. Under the ‘Entrapment-based FDP 1%’ filter, ionbot detected non-canonical peptides in most of the samples, while classic pipelines yield no identifications in 22 out of 24 samples under FDP 1%.

As was noted in the introduction, considering identification properties in local FDR estimation may improve the FDP control. We investigated the influence of filtering strategies on search results (Methods). Peptides were grouped by protein type (canonical/non-canonical) and filtered with group-wise and group-walk algorithms (Figure 1B). Group-based FDR achieves substantially better results for non-canonical peptides for all pipelines, with a visible advantage offered by group-walk filtering. However, the upper bound of group-walk filtering still indicates the FDR control to be suboptimal. In closed search, none of the TPP-run samples meet group-walk FDR 1% threshold, and both FragPipe and ionbot in open search FDP exceed 1% (albeit quite differently, at 57.8% and 3.8% maximum, respectively). Group-based FDR also underestimates FDP for peptides of canonical proteins compared to global FDR, possibly due to confounding groups based on the physical properties of numerous peptides.

Filtering peptide groups based on the estimated FDP rather than the FDR predictably equalizes the FDP across all samples, peptide types, and pipelines to ∼1%. Thus, we enforced an FDP-calibrated filtering to ensure fair comparison of pipeline performance in non-entrapment search. For further analysis, non-entrapment results were filtered on peptide level using thresholds mapped from each sample’s entrapment results. The maximum score rank corresponding to an FDP ≤ 1% of each peptide type in the entrapment search was applied as the threshold. The same approach used to map 1% global FDR shows strong correlation between score ranks in entrapment and non-entrapment searches, suggesting the robustness of the approach (Supplementary Figure S1). Entrapment-based filtering typically yields a smaller subset of identifications compared to global and group FDR approaches when applied to non-canonical peptides (Supplementary Figure S2). Therefore, entrapment-based filtering can be perceived as a more stringent cutoff for low-likelihood detections.

### Pipeline and database affect spectra interpretation

To evaluate the performance of closed and open modification searches across different search space sizes, we compared the output of the Transproteomic Pipeline (TPP) (closed search) and FragPipe (open search) with the corresponding modes of ionbot. To ensure baseline comparability, we harmonized digestion parameters and filtering thresholds across the three tools, leaving other settings at default (Methods). To account for the bias of current LC-MS/MS technologies against non-canonical proteins, we considered a non-canonical protein identified if at least one unique (proteotypic) peptide was detected.

### Ionbot reproducibly identifies non-canonical proteins in closed search mode

TPP and ionbot in closed search mode annotated similar sets of spectra with a substantial overlap (73.9%) (Supplementary Figure S3A). Neither the choice of pipeline nor database size influenced the identification rate (two-way ranked ANOVA: pipeline p=0.43, database p=0.18) (Figure 2A). The identification rate in ionbot is significantly higher than in TPP within only the TrEMBL database search (Wilcoxon p<0.001); however, the difference does not exceed 1.7% per sample on average (Table 1). Ionbot identifies 5% more spectra matching non-canonical peptides than TPP (Figure 2B), which are also found more often in two or more samples than in TPP (Figure 2C-D). Therefore, ionbot offers more robust evidence.

**Figure 2:**
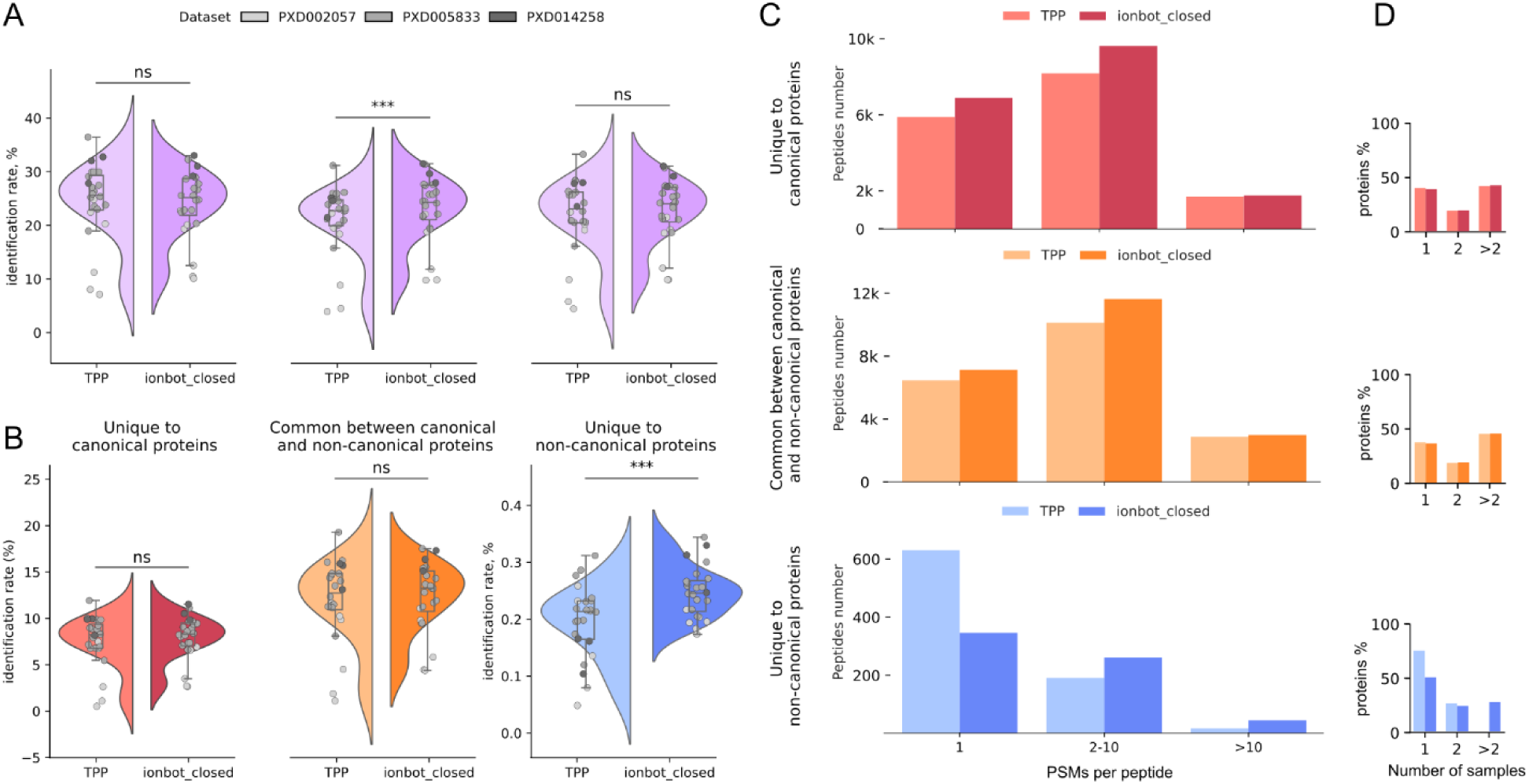
Characterisation of TPP and ionbot closed search results on PSM and peptide level. **A.** Comparison of identification rates (calculated PSM number over spectra number for each sample) of TPP and ionbot across different databases. PSM level was filtered with global FDR of 1%. **B.** Identification rates per peptide type and pipeline from OpenProt search. **C.** Number of PSMs per non-redundant peptides set across all samples from OpenProt search. Peptide level was filtered according to entrapment FDP 1% per protein type (canonical/non-canonical). TPP identifies >1 PSM for 24.6% of non-canonical peptides, while for ionbot this number is 46.8%. **D.** Percent of peptides found in one, two, and more samples across all datasets from OpenProt search, divided by peptide type and pipeline. TPP presents zero non-canonical peptides found in more than two samples.

**Table 1.** Summary of the comparison between pipelines under entrapment-based filtering.

| Search | Closed |  | Open |  |
| --- | --- | --- | --- | --- |
| Pipeline | TPP | ionbot | FragPipe | ionbot |
| Average ID-rate across databases | 20.3% | 21.2% | 29.3% | 36.3% |
| Average ID-rate of non-canonical peptides | 0.19% | 0.25% | 0.81% | 0.94% |
| Average % of reproducible non-canonical peptides (>1 PSMs) | 24.6% | 46.8% | 23.4% | 40.0% |
| Percentage of non-canonical peptides identified within >1 sample | 25.9% | 50.3% | 6.02% | 46.8% |
| Average loss in identified proteins by ionbot compared to classical pipelines across databases | - | 20.7% | - | 11.9% |
| Average percent of unambiguous proteins found >1 samples across databases | 60.7% | 68.6% | 58.8% | 66.9% |
| Percentage of non-canonical proteins supported by >1 PSMs across all samples | 90.9% | 97.2% | 86.8% | 94.7% |
| Percentage of non-canonical proteins supported by >1 peptides across all samples | 68.2% | 65.1% | 52.6% | 65.0% |
| Mean coverage of unique protein sequence by detected peptides (canonical proteins) | 0.284 | 0.305 | 0.353 | 0.414 |
| Mean coverage of unique protein sequence by detected peptides (non-canonical proteins) | 0.284 | 0.227 | 0.346 | 0.305 |
| Percent of proteins < 100 amino acids long (canonical proteins) | 9.8% | 6.8% | 9.6% | 7.2% |
| Percent of proteins <100 amino acids long<br>(non-canonical proteins) | 50.0% | 41.3% | 42.1% | 42.1% |

“-” means that the comparison is not relevant.

TPP outperforms in protein identifications for all three databases (Figure 3A). Nevertheless, the detection frequency across samples is similar between pipelines for both unambiguous and indistinguishable proteins (Figure 3B). However, ionbot uncovers ∼five times more non-canonical proteins with proteotypic peptides supported by multiple PSMs compared to TPP (Figure 3C-D). This is relevant, as the large number of protein sequences in OpenProt makes it statistically unlikely that two random (i.e., erroneous) peptides would match the same protein. The TPP’s proteotypic peptides show the opposite pattern, where these peptides cover a larger fraction of unique sequences in non-canonical proteins than ionbot’s, and favor shorter proteins (50% are shorter than 100 aa) (Figure 3E). This may indicate a more random scattering of identified peptides across the entire database and deviates from the overall TPP coverage distribution for canonical and common peptides (Figure 2D). This deviation is much less prominent in ionbot results. When evaluated at the peptide level, however, both pipelines yield comparable results, where TPP is more conservative under entrapment-based filtering to non-canonical proteins than ionbot in closed search.

**Figure 3:**
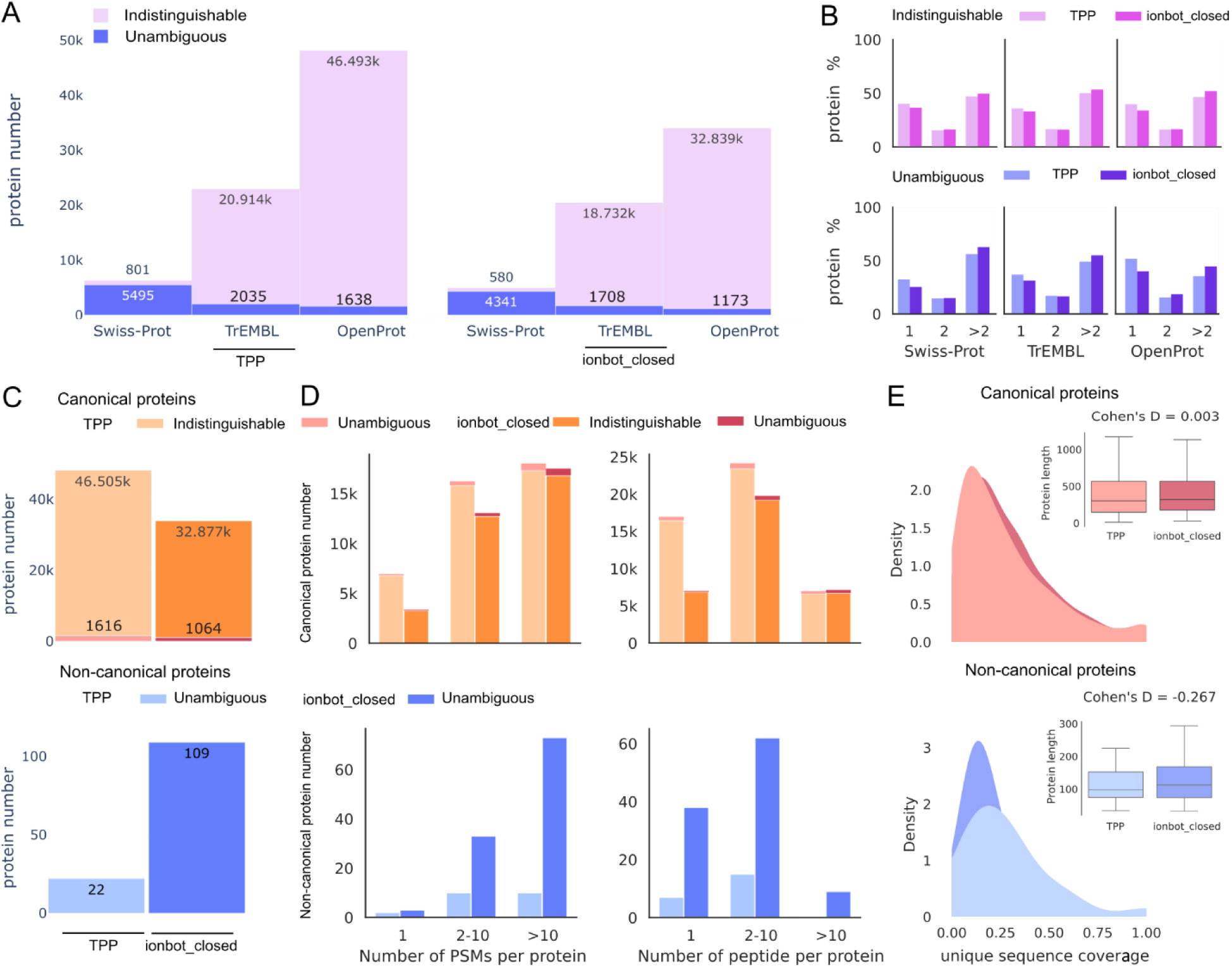
Characterisation of TPP and ionbot in closed mode results on protein level. **A.** Number of identified proteins with each pipeline for all three databases. Proteins identified in several samples were counted once. Proteins identified with at least one unique peptide are referred to as unambiguous, and proteins identified with no unique peptides are referred to as indistinguishable proteins. **B.** Percent of proteins found in one, two, and more than two samples across datasets for each pipeline from OpenProt search. **C.** Number of identifications per protein type using OpenProt between pipelines. Proteins identified in several samples were counted once. Indistinguishable proteins were classified as canonical if at least 1 protein in the protein group was canonical. Non-canonical proteins were considered only if detected with at least one proteotypic peptide. **D.** Number of PSMs (left) and peptides (right) per protein in each protein type and pipeline. Proteins and peptides identified in at least one sample were counted once. **E.** Percentage of residues of unique protein sequence covered by non-redundant peptides from OpenProt search, per protein type and pipeline. Proteotypic peptides were extracted from *in silico* digestion of OpenProt. The total extent of overlapping unique peptides per protein was considered as "unique area". Coverage was calculated as the ratio of amino acids covered by detected proteotypic peptides to the theoretical "unique area" length.

### Ionbot corroborates and improves upon the results of other open modification search engines

FragPipe shares 73.6% of identified spectra with ionbot run in open search mode (Supplementary Figure S3B). In both pipelines, the identification rate largely depends on pipeline choice but not on database size (two-way ranked ANOVA: pipeline p<1.8×10^−8^, database p=0.094) (Figure 4A). Ionbot’s open search consistently identifies more spectra than FragPipe (Table 1), increasing spectrum-level identifications by 5.6% (on average) across samples (Wilcoxon test p<0.001). Like with TPP, ionbot identifies fewer canonical proteins, but ∼5.5 times more non-canonical proteins under entrapment-based filtering, despite a non-significant gain on identification rate (Figure 4B, Figure 5A, 5C). Whereas protein reproducibility is similar between pipelines (Figure 5B), non-canonical proteins are also better supported by ionbot: 94.7% are backed by multiple PSMs or peptides in more than one sample, compared to 86.8% in FragPipe (Figure 4C, 5D). Despite these differences, both pipelines cover a third of the unique non-canonical protein sequences (Figure 5E).

**Figure 4.**
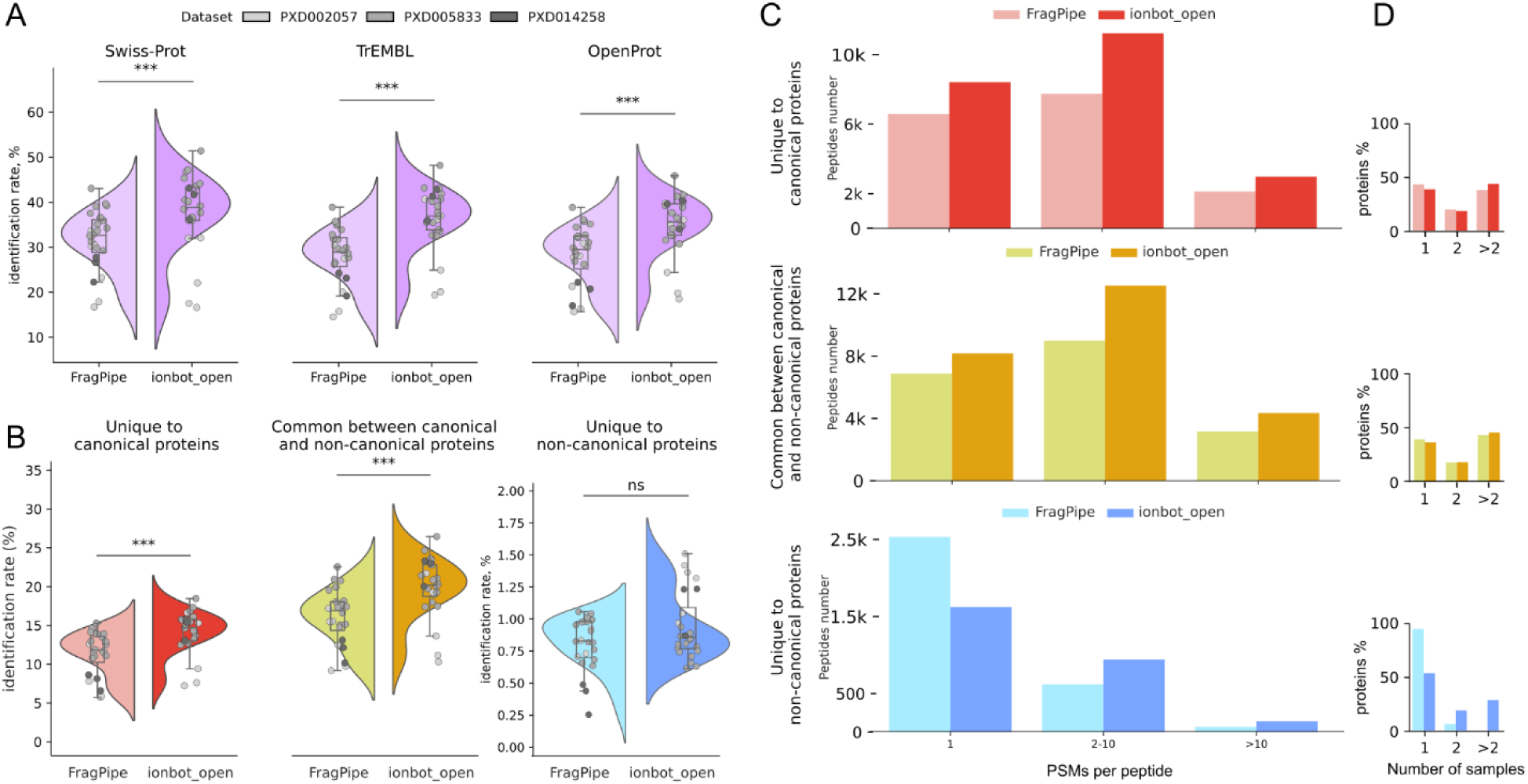
Characterisation of FragPipe and ionbot in open search mode results on PSM and peptide level. **A.** Comparison of identification rates (i.e., number of PSMs over number of spectra for each sample) of FragPipe and ionbot across different databases. Results were filtered to global FDR 1% at the PSM level. **B.** Identification rates per peptide type and pipeline from OpenProt search. **C.** Number of PSMs per non-redundant peptide set across all samples from OpenProt search. Peptide level was filtered according to entrapment FDP 1% per protein type (canonical/non-canonical). **D.** Percent of peptides found in one, two, and more samples across all datasets from OpenProt search, divided by peptide type and pipeline. FragPipe presents zero non-canonical peptides found in more than one sample.

**Figure 5.**
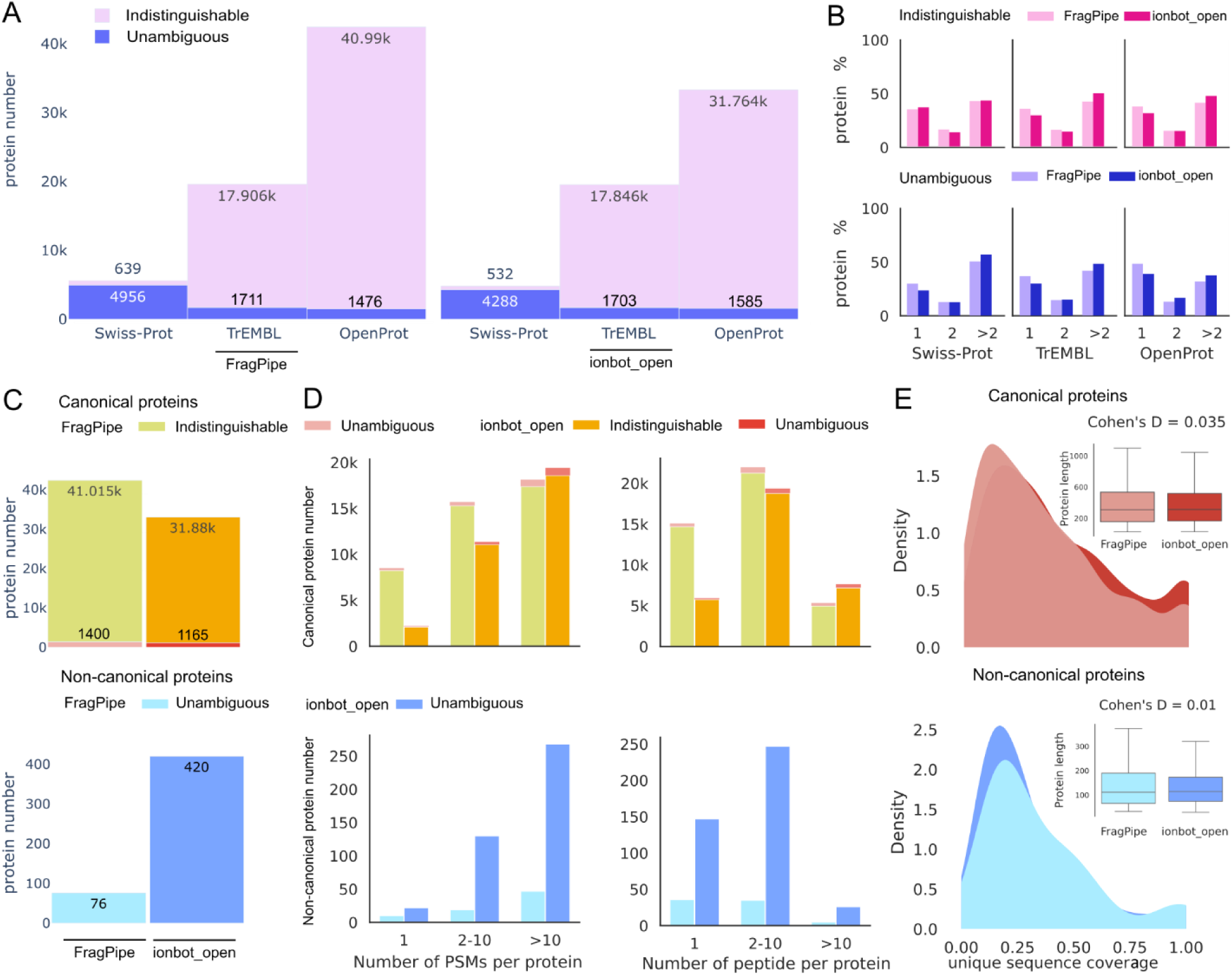
Characterisation of FragPipe and ionbot in open search mode results on protein level. **A.** Number of identified proteins with each pipeline for all three databases. Proteins identified in several samples were counted once. Proteins identified with at least one unique peptide are referred to as unambiguous, and proteins identified with no unique peptide are referred to as indistinguishable proteins. **B.** Percent of proteins found in one, two, and more than two samples (across datasets) for each pipeline from OpenProt search. **C.** Number of identifications per protein type using OpenProt between pipelines. Proteins identified in several samples were counted once. Indistinguishable proteins were classified as canonical if at least one protein in the protein group was canonical. Non-canonical proteins were considered only if detected with at least one proteotypic peptide. **D.** Number of PSMs (left) and peptides (right) per protein in each protein type and pipeline. Proteins and peptides identified in at least one sample were counted once. **E.** Percent of residues of unique protein sequence covered by identified peptides per protein type and pipeline from OpenProt search.

Differences in peptide-to-spectrum matching also influence how PTMs are annotated. FragPipe annotates up to two known mass shifts per PSM, while ionbot can only deconvolute one unexpected mass shift (selected from Unimod repository). While 67.2% of commonly identified spectra are annotated similarly (both pipelines annotate the spectra with unmodified/variable or unexpected mass-shift peptides), over 32.8% show divergent interpretations between pipelines (Supplementary Figure S3C). This highlights how scoring models and search strategies influence not just sensitivity, but also biological conclusions drawn from PTM patterns.

### Comprehensive database reassigns false spectrum matches to non-canonical proteins

Peptide-spectrum interpretation is inherently search-space dependent. Larger databases introduce additional sequences, which may better explain observed spectra, potentially rescuing spectra that remain unidentified in smaller databases. Paradoxically, expanding the search space can reduce the number of identifications at global FDR<1%, due to the increased likelihood of matches to high-scoring decoy sequences. In our study, the TrEMBL database only includes canonical proteins and is smaller than the OpenProt database, which also includes non-canonical proteins. When comparing both searches with ionbot open mode, the vast majority (∼90%) of spectra were identified as canonical peptides in both searches (Figure 6A; Supplementary Table S1). However, 1.4% were reannotated as non-canonical, highlighting the influence of the database on spectrum interpretation. Some spectra that had remained unidentified in the narrower search were matched to canonical (0.9%) or non-canonical (1.1%) sequences, highlighting a gain from reinterpreting previously unassigned signals. At the same time, several confident identifications in TrEMBL no longer passed the FDR filter in OpenProt (5.9%), reflecting the trade-off introduced by a larger search space. Overall, non-canonical identifications in OpenProt stem from both reannotated spectra and previously unidentified ones (Figure 6B).

**Figure 6.**
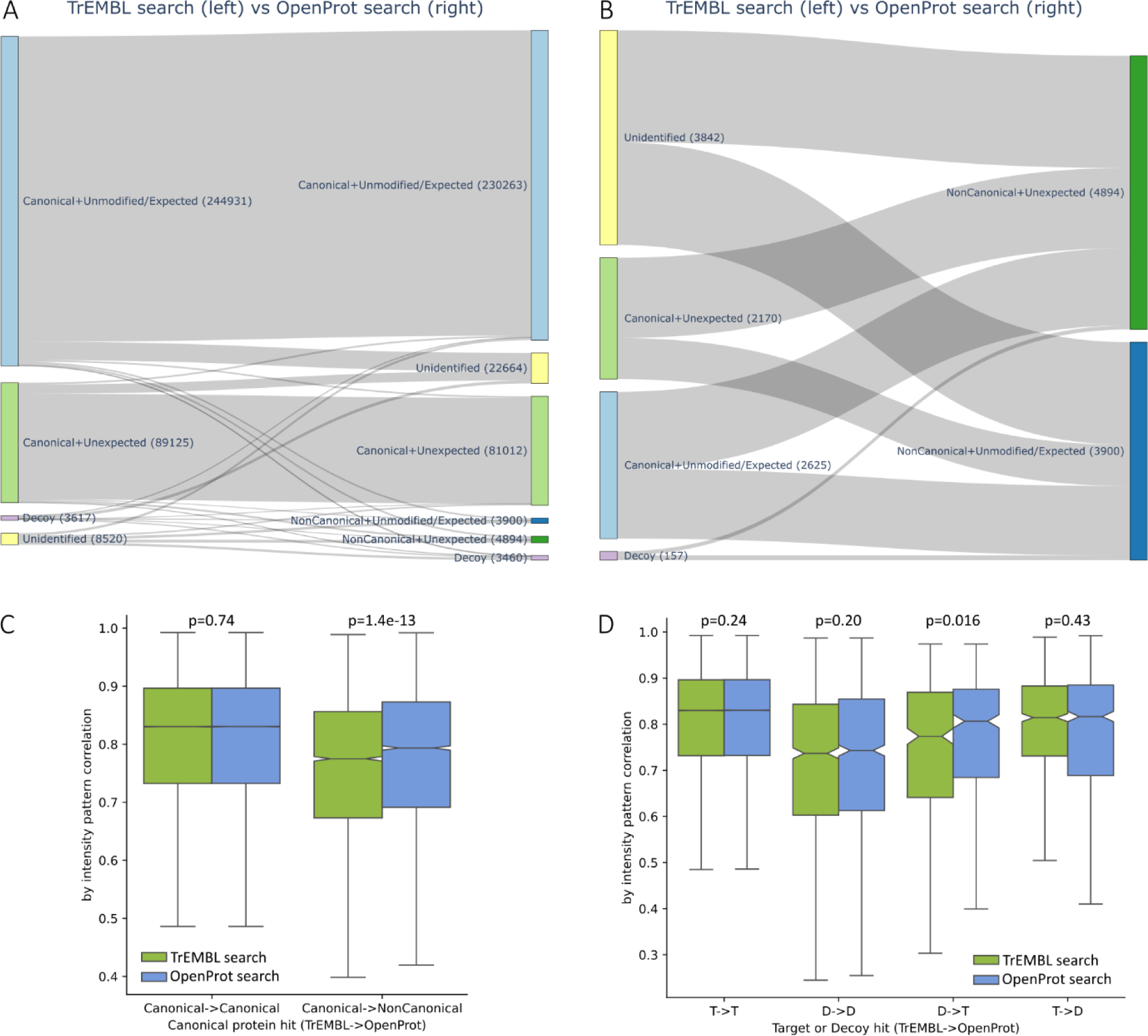
Comparison between ionbot searches against TrEMBL and OpenProt. **A.** Sankey plot showing how some spectra are interpreted differently when conducting an open search against TrEMBL (left) and against OpenProt (right). The plot shows target and decoy PSMs with q-value < 0.01 (global FDR filtering). Spectra that were identified (or passed the q-value filtering) only in one of the two searches were labelled “Unidentified” in the other. **B.** Zoomed Sankey plot, highlighting the source of non-canonical peptides identifications. Although most non-canonical peptides derive from spectra unidentified in the open search (rescued spectra), a considerable amount derive from re-annotation of false positive hits in the TrEMBL search. In addition, non-canonical peptides with unexpected modifications are more common than those without. **C.** Comparison of by-ion intensity correlation for spectra matched to canonical or non-canonical peptides in the two searches. **D.** Comparison of by-ion intensity correlation for spectra matched to target or decoy peptides in the two searches.

Since OpenProt includes TrEMBL sequences, we examined why spectrum annotations changed. We compared the by-ion intensity correlation for matching peptides (Figure 6C). For spectra reannotated from canonical to non-canonical, the OpenProt PSMs consistently showed a stronger agreement between experimental and theoretical spectra, especially for mass shifts beyond variable modifications (p=2.7×10^-16^, Mann-Whitney U test). This indicates that these new matches are not only different but also better supported by the experimental evidence and thus provide a solid basis for downstream analysis.

To ensure this improvement was not merely due to the increased search space, we compared the by-ion intensity correlations of decoy matches across both searches. Decoy identifications in the OpenProt search do not show significantly higher intensity correlations compared to the corresponding identifications in the TrEMBL search (Figure 6D). Therefore, we concluded that the difference in correlation shown in Figure 6C can be attributed to the higher quality of the peptide identifications with the OpenProt database.

### Open search can correct false-positive non-canonical peptide identifications

Open modification search also increases the search space considerably. We compared the results obtained with ionbot in closed search mode with its results in open search mode for the OpenProt database to investigate the impact of open search. Ionbot open modification search increased the overall identification rate by 60% compared to closed search (Figure 7A). Of the new PSMs, 4.3% were assigned exclusively to non-canonical proteins (Supplementary Table S1, Figure 7B). We found that 9.8% of the spectra identified in the closed search could be explained by a different peptide bearing an unexpected modification in the open search. More specifically, the open search reassigned 38.1% of non-canonical PSMs from the closed search to canonical peptides bearing an unexpected modification, while only 1% of closed search canonical PSMs were reassigned as non-canonical peptides bearing an unexpected modification. On the peptide level, the open search identified 817 additional proteotypic peptides and tripled the number of non-canonical proteins detected to 697. The number of non-canonical proteins supported by two or more peptides rose six-fold (from 31 in closed search, to 184 in open search), despite the share of non-canonical peptides supported by two or more PSMs being comparable. Furthermore, the portion of non-canonical peptides found in three or more samples in open search dropped by 9.12% on average, possibly due to sample-specific non-canonical proteins.

**Figure 7.**
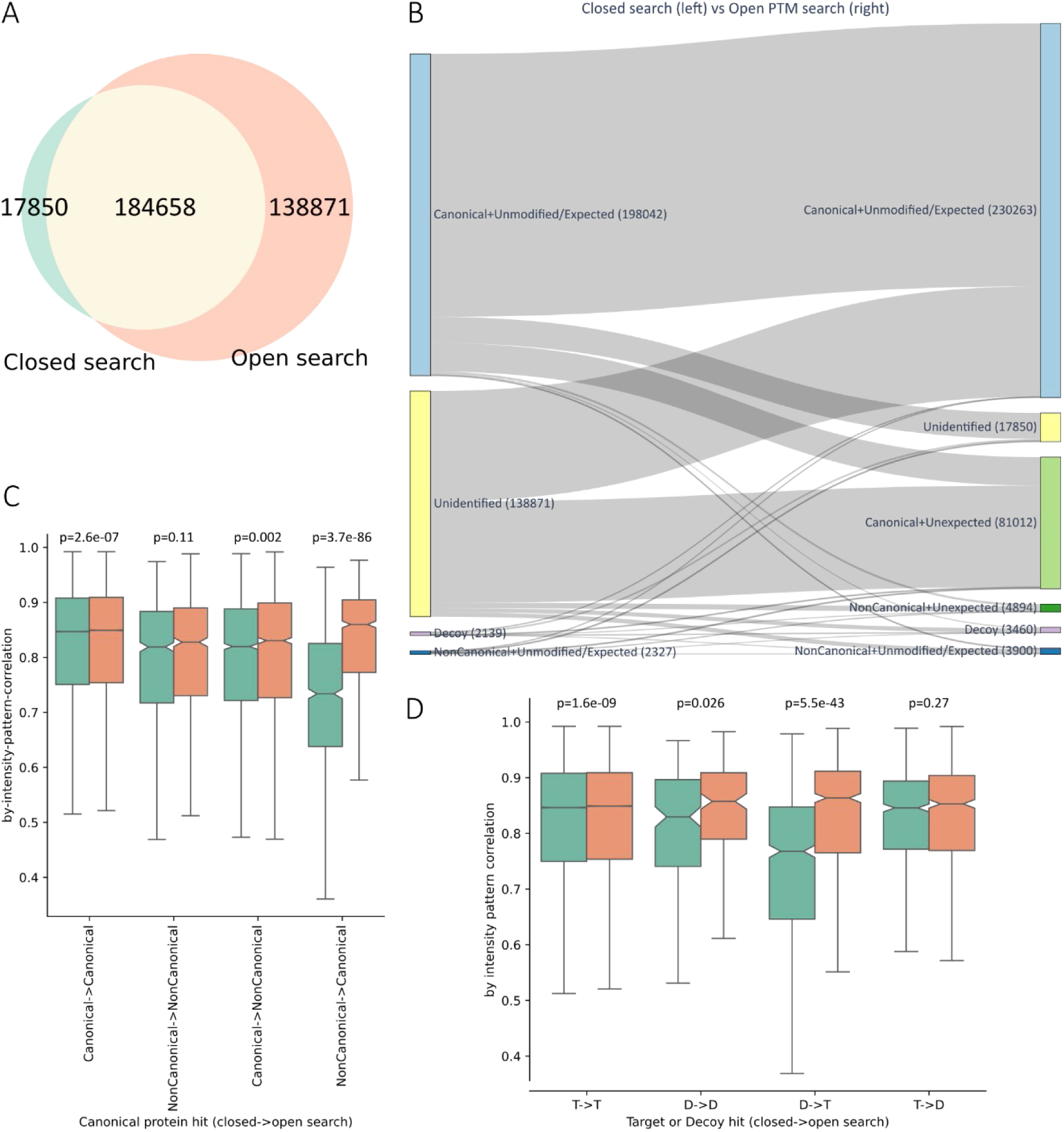
Comparison between ionbot closed search and open search results. **A.** Venn diagram showing the spectra identified by the two searches. The open search also identified nearly all spectra that were identified in the closed search. **B.** Sankey plot showing how some spectra are interpreted differently when conducting a closed search (left) and an open search (right) against the OpenProt database. The plot shows target and decoy PSMs with q-value < 0.01. Spectra that were identified (or passed the q-value filtering) in only one of the two searches were labelled “Unidentified” in the other. Over 130,000 spectra unidentified in the closed search were rescued by the open search, leading to a considerable increase in the overall identification rate. **C.** Comparison of by-ion intensity correlation for spectra matched to peptides with or without unexpected modifications in the two searches. **D.** Comparison of by-ion intensity correlation for spectra matched to target or decoy peptides in the two searches.

We once again investigated the PSMs in the two searches to ensure that spectra reannotated in the open search have higher by-ion intensity correlation (Figure 7C). In this comparison, the intensity correlation improvement is much larger than what we observe for the decoys (Figure 7D), strongly suggesting this improvement is not an artefact derived from the increased search space. Overall, ionbot open search provides clear quantitative and qualitative gains for non-canonical protein identification.

### Open search reveals characteristic modification patterns of non-canonical proteins

Since 17.7% of mass shifts can correspond to multiple PTMs, we grouped PTMs by mass shifts. Surprisingly, non-canonical peptides show different mass shifts compared to canonical and common peptides. Non-canonical proteins have a higher proportion of modified-to-unmodified peptides, underscoring their reliance on open search for identification (Figure 8A). Furthermore, 71.7% of modified non-canonical peptides exhibited mass shifts included unexpected modifications, compared to 34.7% among canonical peptides. This results in 90.6% of identified non-canonical proteins being modified at least once, versus 68.9% of canonical proteins (Figure 8B). Importantly, 11.5% of proteins represented solely by modified peptides are non-canonical (Figure 8C), making them undetectable in closed searches. Thus, these results align with our hypothesis that open modification search in proteogenomics allows us to detect novel proteins more effectively.

**Figure 8:**
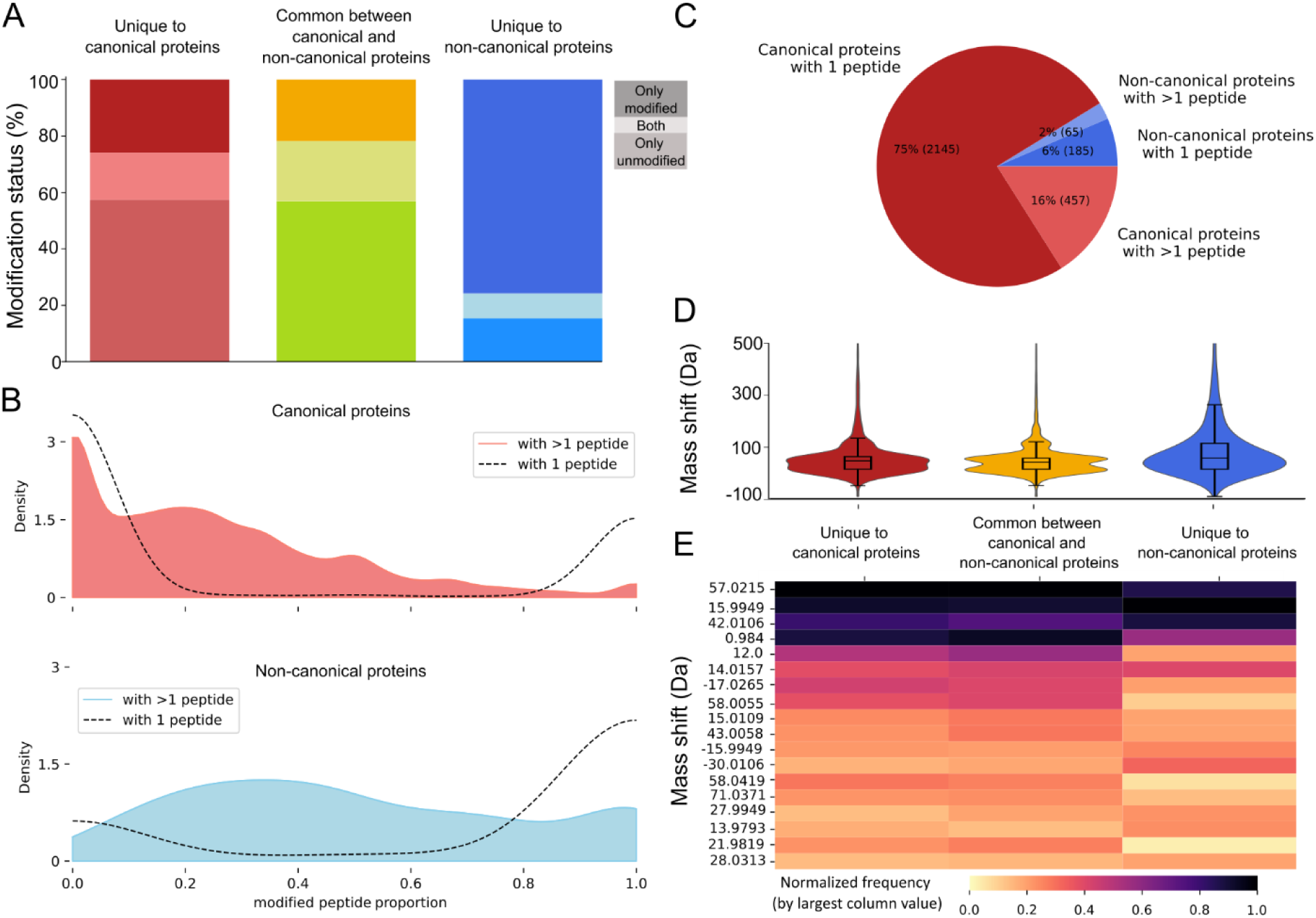
Importance of open modification search. **A.** The proportion of peptides detected always with modification (top of bar), both modified and unmodified (middle of bar), and never modified per peptide type (bottom of bar). **B.** Frequency distribution of modified peptides in canonical and non-canonical proteins. The colored area shows proteins with more than one detected peptide. Dashed lines show proteins detected with only one peptide (18.5% canonical and 75.6% non-canonical proteins). Canonical proteins with low modification rates are found more often (mean 0.367). Single-peptide canonical proteins have less probability of being modified (27.3%) than unmodified. Non-canonical proteins detected with more than one peptide are modified more often than canonical (mean 0.908). 88.5% of non-canonical proteins with single-peptide detection are only modified. **C.** Pie chart of proteins whose identified peptides are all modified. Canonical proteins constitute a considerable fraction of modified proteins (88.5%) suggesting that, like non-canonical proteins, many canonical proteins are overlooked in closed search. **D.** Distribution of mass shifts of modified peptides. The range is cropped at 500 Da for easy visualisation. **E.** Heatmap showing the fifteen most frequent mass shifts in the datasets for each peptide type (excluding mass shift of zero – i.e., unmodified peptides). The frequencies were normalized by largest column value. Mass shifts corresponding to common artefacts like carbamidomethylation (+57.0215 Da), M-oxidation (+15.9949 Da), and deamidation (+0.9840 Da) appear with high frequency across all three peptide categories, in line with expectations. Other PTMs show different patterns in non-canonical peptides compared to canonical and common peptides.

Mass shift distributions further illustrate these differences. Common peptides show two prominent peaks at +0.9840 Da and +57.0215 Da (Figure 8D). The +0.9840 Da mass shift can reflect amino acid substitutions (Asparagine ◊ Aspartic acid, Glutamine ◊ Glutamic acid) or deamidation, the latter arising both physiologically and as an LC-MS/MS artefact [36,37]. The +57.0215 Da shift corresponds mainly to carbamidomethylation, a sample preparation artefact, but can also be attributable to two amino acid substitutions (Alanine ◊ Glutamine, Glycine ◊ Asparagine). Together with two other most frequent mass shifts (N-terminal acetylation (+42.106 Da) and oxidation (+15.9949 Da)), these modifications vary in relative abundance among peptide types (Figure 8E). Non-canonical peptides are depleted in unexpected shifts at +58.0055 Da, +58.0419 Da, +71.0371 Da, and enriched for +13.9793 Da, +14.9997 Da, +27.0473 Da compared to canonical peptides (>=0.05 absolute difference, Figure 8E). These mass shifts are either chemical derivatives or signs of amino acid substitution [38]. It is expected that sample preparation artefacts are prevalent among mass shifts. However, we also identified phosphorylation and ubiquitination – both well-known biological PTMs – on one in seven proteotypic non-canonical peptides. Together, these patterns highlight unique modification landscapes in non-canonical proteins and emphasize the value of open modification search in revealing these.

### Analysis of modifications isobaric to single amino acid substitutions identifies potential protein variants

Among the twenty most frequently detected modifications, fifteen correspond to mass shifts isobaric to amino acid substitutions. These mass shifts may indicate single amino acid variants (SAAVs), but distinguishing PTMs from SAAVs is challenging without matched genomic data or non-cancer proteomes available. Because OpenProt was built by predicting non-canonical ORFs from the reference genome, mutations are not included, and we might identify mutated versions of canonical peptides that map to non-canonical proteins by chance, or variants of canonical and non-canonical proteins absent in OpenProt. One way to handle isoSAAVs is to inspect if a corresponding genomic region has ever been observed with an expected missense mutation. As a source of global genomic variation, we utilized gnomAD, a database of more than 200 million small genomic variants of ∼150,000 exons and genomes from Eurasian, African, and American populations [39].

We found amino acid substitutions isobaric to mass shifts (isoSAAVs) in 41.3% of proteotypic peptides from canonical proteins (1,312 peptides, 800 proteins), and in 78.8% of proteotypic peptides from non-canonical proteins (416 peptides, 337 proteins). Analysing isoSAAVs in peptides shared between multiple proteins is more challenging due to their ambiguous genomic origin. 46.6% (1,728) of shared peptides were detected with at least one PSM with isoSAAV. These isoSAAVs were cross-referenced with 131,963 gnomAD positions, 57% of which contained known single nucleotide variants (SNV), with similar number of affected non-canonical (823) and canonical (772) proteotypic peptides. Shared peptides contained 17,277 known SNVs collected from all possible origins of peptides. Of the inspected isoSAAV positions, 7,323 led to expected amino acid substitutions, with 4.7% variants in 212 proteotypic canonical peptides (153 proteins), 5.9% variants in 212 proteotypic non-canonical peptides (186 proteins), and 89.4% variants in 3,916 shared peptides.

Thus, out of 13,941 PSMs with a mass shift (entrapment-based selected peptides), 3% of proteotypic (1.6% non-canonical and 1.4% canonical PSMs) and 31.7% of shared PSMs could be explained by an SAAV rather than a modification. Our analysis thus presents a framework and highlights the importance of verifying open modification search results against genomic variation data.

### Search parameters determine what peptides we see

Open search strategies expand the detectable peptide space by incorporating mass shifts. Consequently, ionbot open search yields an additional 32.5–37.9% long peptides (longer than fifteen amino acids) for both canonical and non-canonical peptides relative to closed search (Figure 9A). Because longer peptides exhibit a higher likelihood of being proteotypic, this increase contributes to a higher rate of confident protein identifications. Accordingly, the amino acid composition of detected peptides is expected to change between different search spaces. 91.7% of the mass shifts of PTMs are specific to 1-3 residues. For example, phosphorylation is inherent to Serine, Threonine, and Tyrosine. Non-canonical proteins have been shown to have a distinct amino acid composition, which has a small effect on their detectability with LC-MS/MS and closed search [5,41], and which can also affect the likelihood of a peptide bearing specific modifications. We examined how the amino acid frequency of detected peptides changes between search type (closed vs open), and peptide type (canonical vs non-canonical), which revealed that discrepancies in amino acid composition between canonical and non-canonical peptides did not depend on search type; however, the relative abundance of several residues was affected by search parameters (Figure 9, left). Notably, these differing residues correspond to three or more of the most frequent mass shifts detected in open search (Figure 8E), underscoring the association between detected peptides and modifications. Canonical and non-canonical proteins with mass spectrometry evidence in OpenProt have a significantly different frequency for seven amino acids (Figure 9, right). Although 75% and 85% of residues showed a negligible difference between types of peptides for open and closed searches (|Cohen’s D|<=0.2), a few amino acids were either depleted in canonical peptides compared to non-canonical (Asparagine in open search, Isoleucine and Methionine in both searches), or vice versa (Proline in open search and Arginine in both searches), highlighting a small contribution of search parameters to detection bias (Figure 9, right). Interestingly, Methionine is overrepresented in non-canonical peptides compared to canonical peptides in both searches and an *in silico* digested database, owing to a higher proportion of N-terminal peptides detected (Figure 9C). In theory, 35.6% of canonical and 86.1% of non-canonical proteins produce a proteotypic N-terminal peptide of detectable length after trypsin digestion. However, ionbot open searches with entrapment-based filtering yield a comparable number of canonical and non-canonical proteins detected by an N-terminal peptide (Figure 9C). The decomposed modification pattern illustrates the positional bias (Figure 9E); half of the mass shift differences between proteotypic peptide types are carried by internal peptides, and the other half by N-terminal peptides (>=0.05 absolute difference, Figure 9E).

**Figure 9.**
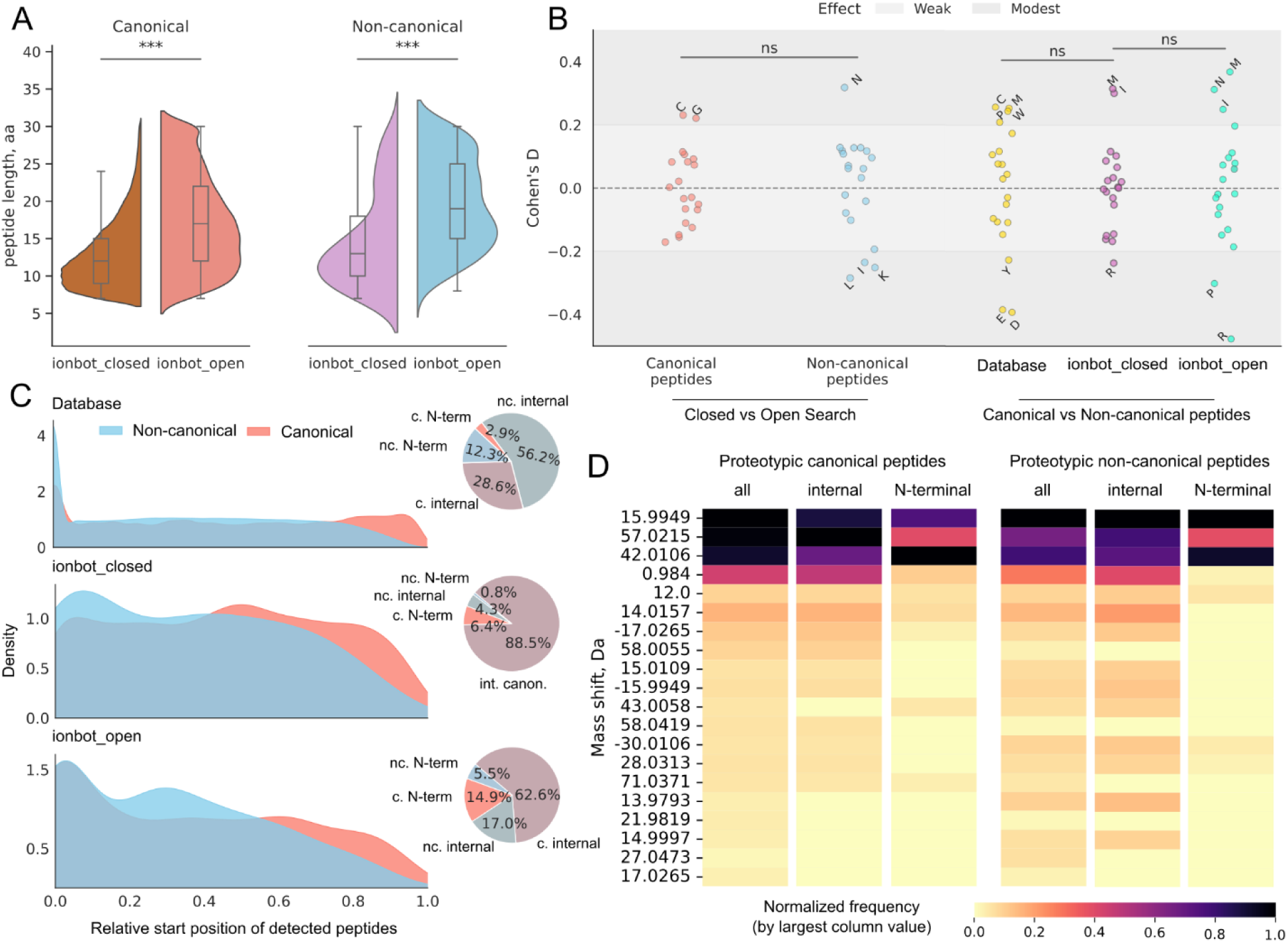
The influence of search parameters on observed peptides characteristics. **A**. Peptide length depending on peptide type and source. Detected peptides are significantly longer in open search than in closed search (Mann Whitney p<0.001). **B.** Amino acid composition of peptides depending on their type and source. Each dot represents the difference in amino acid frequency between two categories, expressed as Cohen’s D statistic. Left: Amino acid composition difference between detected peptides of the same type in different search spaces. Cysteine (C) and Glycine (G) were modestly overrepresented in open search compared to closed search for canonical peptides (Cohen’s D>0.2). The non-canonical peptides in open search were enriched in Asparagine (N) and depleted in Leucine (L), Isoleucine (I) and Lysin (K) in comparison to closed search. Right: comparison of peptides from *in silico* digested proteins with proteomic evidence registered by OpenProt (in yellow), comparison of detected peptides by ionbot closed search (in violet), and open search (in cyan). A higher Cohen’s D value indicates that the mean of the open/non-canonical distribution (left/right) exceeds that of the closed/canonical distribution. The differences in effect size between peptide types (left) and peptide sources (right) are non-significant (ns). Despite this, amino acid set with |effect size| > 0.2 (labelled) vary between comparisons. For example, on the right, the effect size becomes more prominent for Arginine (R) and less prominent for Isoleucine (I) in open search compared to closed search. **C.** Distributions of relative start position of proteotypic peptides in proteins from an *in silico* digested database (only proteins with mass-spectrometry evidence in OpenProt) and proteotypic detections in closed and open searches. The open search more closely preserves the positional distribution of database peptides than the closed search. The peptides mapping to the protein N-terminus (with and without Methionine) are considered N-terminal while all others are considered internal. The relative abundance of peptides by position and type is summarised in pie charts (c. = canonical, nc. = non-canonical). Percent of proteins detectable by proteotypic N-terminal peptide (excluding protein groups): 12.9% canonical and 18.3% non-canonical in closed search, 22.1% canonical and 25.5% non-canonical in open search. In the open search, N-terminal to internal peptides ratio is 0.23:1 for canonical and 0.32:1 for non-canonical types, underscoring a slight prevalence of N-terminal peptides among non-canonical proteins. **D.** Top frequent modifications of prototypic canonical and non-canonical peptides in bulk and separated by position. Modifications are ordered as in Figure 8E. Peptide position contributes to differences in peptide type modification profiles.

Therefore, our analysis dissected the correlation behind detected peptides and frequent mass shifts with open search; however, elucidation of the causality of this link and its potential biological relevance will require more in-depth analysis.

In theory, 92% of canonical and 93% of non-canonical proteins produce an N-terminal peptide of detectable length after trypsin digestion. However, non-canonical proteins are more likely to be detected through an N-terminal peptide (17.7% of theoretical peptides unique to non-canonical proteins) than canonical (3.8%) (Figure 10A). Our results with ionbot open modification search follow these expectations for peptides unique to canonical proteins (4.1% peptides are N-terminal), but not for non-canonical proteins (Figure 10A). In open search, 28.0% of detected non-canonical peptides are located on N-termini. Furthermore, there is an increase in N-terminal non-canonical peptides in open search compared to closed search (26.6%), though it is not significant (Chi2 p=0.18). As expected, non-canonical N-terminal peptides have a significantly higher frequency of Methionine than internal peptides (Cohen’s D=0.68). However, this is not the case for canonical N-terminal peptides (Figure 10B). Our analysis included modifications that could capture N-terminal peptides, such as variable oxidation of Methionine (+15.9949 Da) and N-terminal acetylation (+42.0106 Da). Indeed, these modifications seem to be a driving force behind the higher detectability of non-canonical N-terminal peptides while being less frequent on internal peptides (Chi2 p1.5×10^-15^). Interestingly, the profiles of top mass shifts for canonical and non-canonical peptides are closer for N-terminal (Spearman’s rank correlation 0.75) than for internal peptides (-0.17) (Figure 10C). Thus, the positional bias of detected non-canonical peptides is linked to the variable modifications included in the search space.

**Figure 10:**
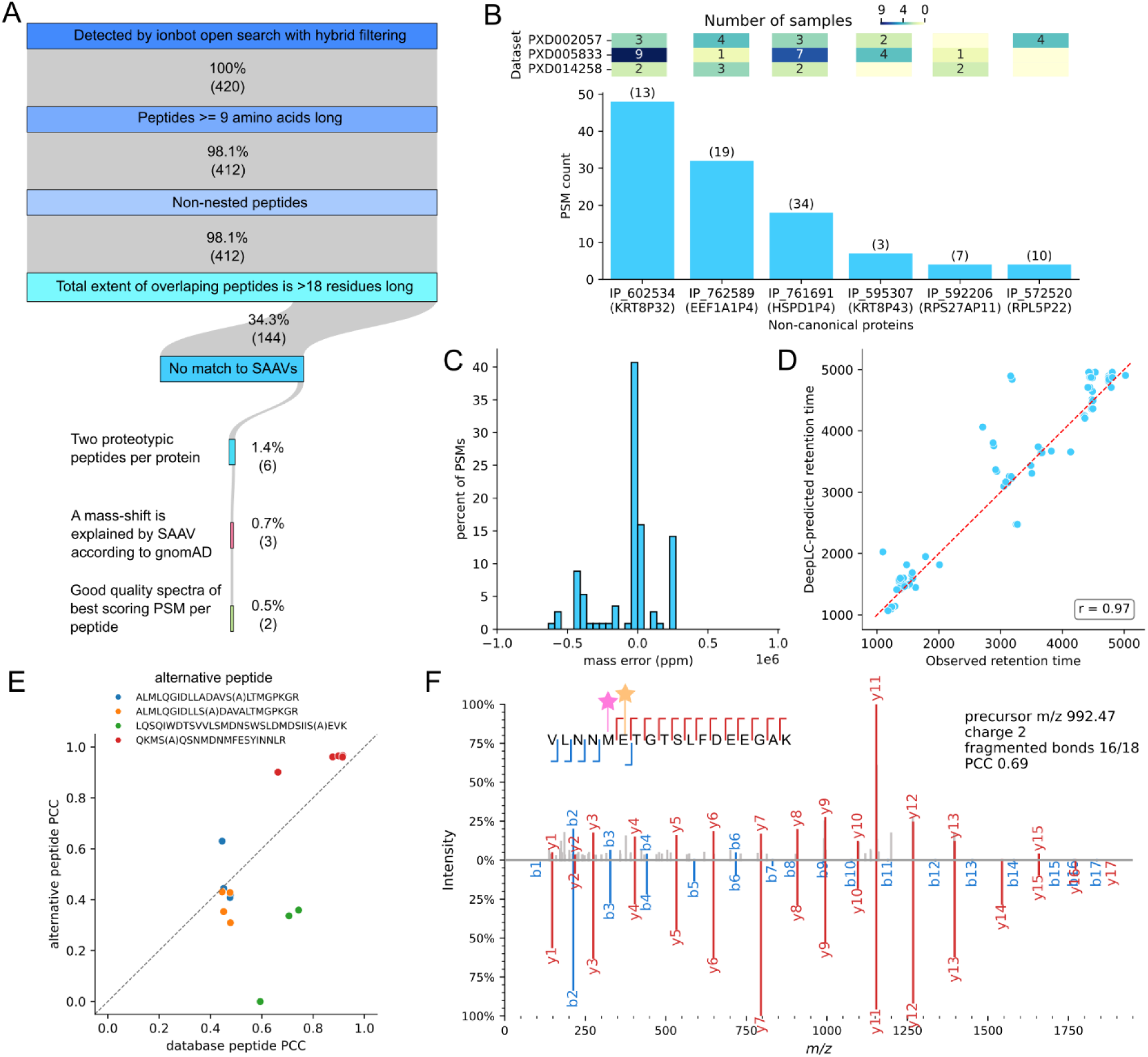
Open-modification search drives novel identifications. **A.** The gradual filtering of non-canonical proteins according to Human Proteome Project (HPP) Guidelines (until step “Two proteotypic peptides per protein”) and spectrum quality. The initial set of non-canonical proteins (n=420) is composed of proteins with at least one proteotypic peptide. The full list of detections on PSM level can be found in Supplementary Table S2 and PSMs can be viewed in Supplementary File S1. The major filtration steps during quality control were the discarding of non-canonical peptides matching known SAAVs of canonical proteins and the two proteotypic peptides requirement. The step “A mass shift is explained by a SAAV according to gnomAD” included filtering out proteins supported by only a single proteotypic peptide, whereas the subsequent step did not apply this restriction. **B.** Top: Number of samples per dataset in which each of the proteins’ peptides were detected. Five proteins were detected in more than one dataset. Bottom: Number of PSMs supporting identification of six non-canonical proteins (all are products of pseudogenes) that pass HPP guidelines. Numbers above bars indicate mass spectrometry-based detection in OpenProt (MS score). **C.** Mass error between precursor and matched peptides’ mass in parts per million (ppm) for identifications that pass HPP guidelines. **D.** Correlation between observed and DeepLC-predicted retention time of peptides that pass HPP guidelines. DeepLC predictions are referenced from the ionbot output files. **E.** Comparison of spectra matched to three OpenProt peptides whose observed mass shifts correspond to amino-acid substitutions or gnomAD missense variants originating from HPP-passing non-canonical proteins. These spectra were further evaluated against the corresponding single amino acid variant (SAAV) peptides, including one case with two alternative peptide forms. The reference residue is indicated in brackets to the right of the variant amino acid. Pearson’s correlation coefficient (PCC) between experimental and predicted spectrum intensity is used as the metric for match quality. Values above the diagonal indicate a better match for alternative peptides. The residues substituted in alternative peptides are highlighted for database peptides if at least one spectrum had a higher PCC with the alternative peptide than the original. The detailed information is listed in Table 2. **F.** Example of a high-quality PSM that was nevertheless removed during the “two proteotypic peptides per protein” filtering step. The peptide is proteotypic to the IP_558630 non-canonical protein encoded by Phosphoglycerate Kinase 1 Pseudogene 1 (PGK1P1). The upper plot is the annotated spectrum, and the lower is the MS2PIP-predicted spectra. It is an 84-residue protein with 94.05% homology to positions 227-310 of the PGK1 gene canonical protein (P00558). The detected peptide differs from the canonical sequence for one residue (21T and 258I), and is modified by Oxidation (+15.9949 Da) and addition of methylamine (+13.0316 Da) (indicated by pink and orange stars, respectively). The mass of Threonine (119.0582 Da) with methylamine (132.0898 Da) has a ∼1Da difference from Isoleucine (131.0946 Da), present on the canonical proteins, highlighting the importance of high-resolution instruments in detecting non-canonical peptides. The match was detected with spectrum number 24032 in RAW file AM16.raw, from dataset PXD005833.

**Table 2.**
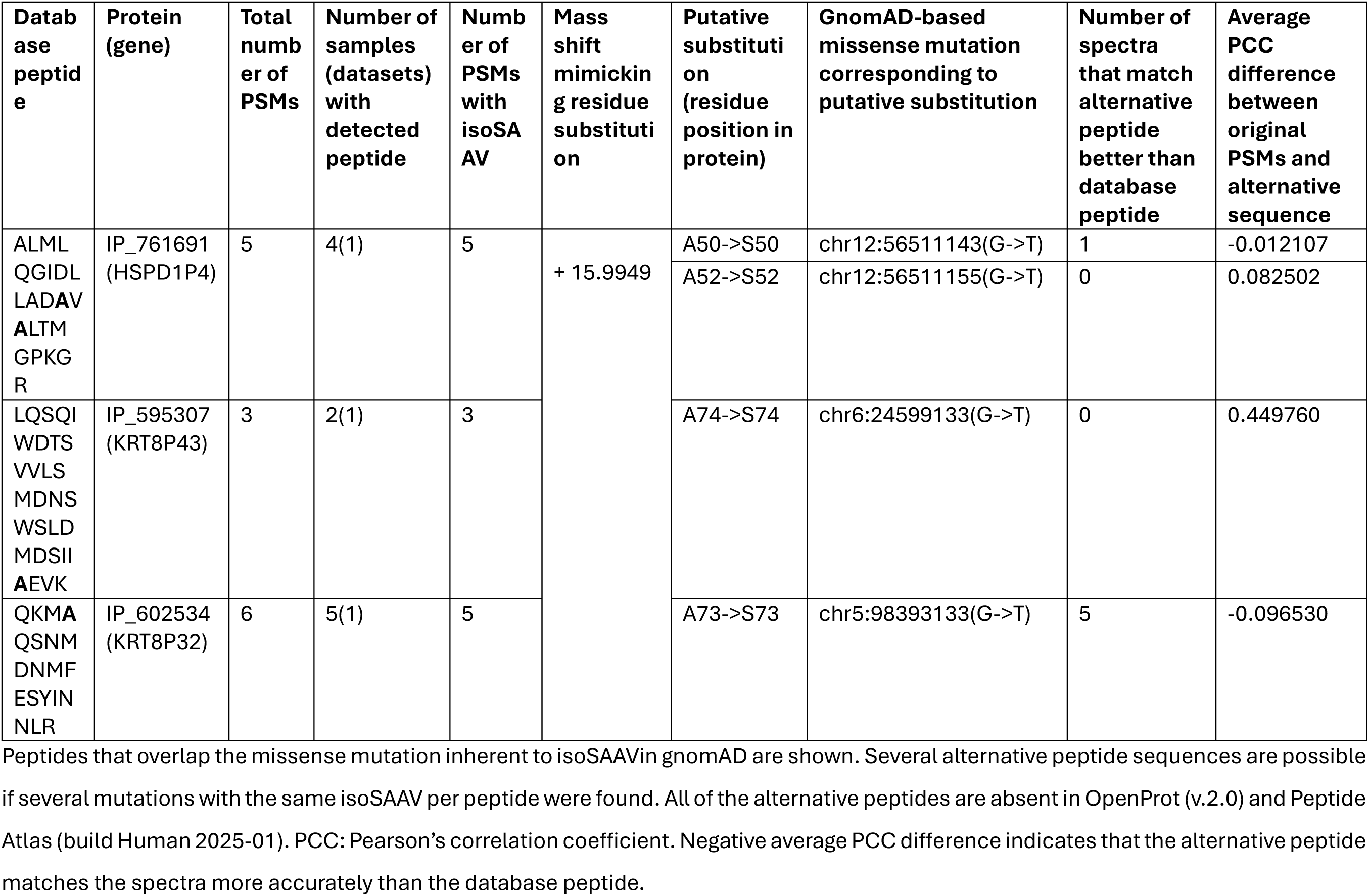
Alternative explanation of proteotypic non-canonical PSMs identified with a mass shift isobaric to re (isoSAAV).

### Rigorous quality control supports modified non-canonical peptides

Proteogenomics inevitably leads to a number of false-positive detections [42]. Therefore, meticulous quality control is necessary in order to claim discovery of a new protein. This would involve providing evidence to classify a protein as PE1 (experimental evidence at the protein level) in accordance with the Human Proteome Project (HPP) data interpretation guidelines. The challenge of providing PE1-level evidence for a non-canonical protein has been discussed before [5]. In theory, 89% (374) of ionbot-detected non-canonical proteins can be detected in compliance with HPP guidelines. However, we could only provide HPP-required evidence for six proteins, as the vast majority of proteotypic non-canonical peptides (516) were either the only peptide mapping to a specific protein, were shorter than nine amino acids, were nested within a longer peptide, or had a total extent with overlapping peptides shorter than eighteen amino acids (Figure 10A). 58% of filtered peptides were mapped to known canonical peptide variants in the HPP portal. The remaining set of non-canonical PSMs exhibits low mass error (|ppm|<1) and strong correlation between observed and predicted retention time (Figure 10C-D).

The six confidently-detected non-canonical proteins have been identified before, and are encoded by open reading frames that overlapped with pseudogenes (Figure 10B). Pseudogenes are copies of protein-coding genes and share high sequence homology with their respective parental genes [43]. 13% of pseudogenic peptides differ from corresponding peptides of parental genes by one or two amino acids [44]. Thus, accurate protein mapping is critical for pseudogenes. Proteotypic peptides of these six non-canonical proteins present at least one PSM with mass-shift isobaric to a single amino acid variant (isoSAAV), which could compromise peptide uniqueness. Of these, three peptides were mapped to known missense mutations in population genomics data (gnomAD), and therefore could be explained by a variant of a canonical protein. To inspect which peptide fits the spectrum better (the pseudogenic peptide or the peptide with SAAV), by-ion peak intensities were compared to predicted intensities of the two sequences by MS2PIP. PCC between predicted and experimental spectra was greater for six of thirteen PSMs with isoSAAV, corresponding to two database peptides (Figure 10E, Table 2). All alternative peptides were absent in OpenProt and Peptide Atlas, putatively featuring a novel variant. All proteins with proteotypic non-canonical peptides with missense mutations found in gnomAD were omitted in further analysis according to HPP guidelines, resulting in three remaining proteins. 66% of proteotypic peptides of confident non-canonical proteins were detected by three or fewer PSM; thus, it is crucial to manually examine PSM quality. We focused on the best-scoring PSMs per peptide and manually validated the quality of six spectra. Two PSMs were marked as passing the assessment, each providing only single-peptide support for two non-canonical proteins (Supplementary Figure S4).

To conclude, the extensive quality control of 420 non-canonical proteins identified by open search led to six proteins complying with the PE1-level evidence required by HPP. However, additional manual spectra examination spotted false positive identifications, illustrating that even recurrent identification of the same protein fragments should be approached with scepticism. An example of a confident spectrum that nevertheless did not satisfy the HPP guidelines is presented in Figure 10F. These findings underscore the necessity of large-scale reanalysis of public datasets to rescue credible one-hit wonders.

All confidently identified non-canonical proteins are encoded by transcripts annotated as pseudogenes in Ensembl (version 106) (Figure 11C). Of the proteotypic peptides corresponding to these non-canonical proteins, 54.5% were detected by only one peptide spectrum match (PSM). Therefore, it is essential to manually examine the quality of these PSMs. We focused on the highest-scoring PSMs for each peptide and manually validated the quality of 83 spectra; only one PSM was classified as high quality (Figure 11F).

In summary, the rigorous quality control of 697 non-canonical proteins identified through open search resulted in 39 proteins meeting the PE1-level evidence requirements set by the HPP. However, further manual examination of the spectra revealed some false positive identifications. The key steps in the quality control process involved discarding non-canonical peptides that matched known SAAVs of canonical proteins and conducting a thorough quality examination of the spectra (Supplementary Figure S2). This highlights that even repeated identification of the same protein fragments should be approached with caution.

## Discussion

Thousands of non-canonical proteins have been detected in humans through a combination of ribosome profiling and novel gene prediction algorithms [45–47]. Yet, experimental and computational limitations still hinder their detection at the peptide level by LC-MS/MS, which remains the gold standard for protein-level identification [5,48]. This work addresses the computational side, focusing on database design and search engine strategy. We evaluate how open modification search strategies can boost the identification of non-canonical proteins. Among the evaluated tools, ionbot showed greater reproducibility compared to TPP, the current gold standard for the identification and validation of new proteins [49], and FragPipe, a popular open search pipeline [24].

Despite the 33-fold increase in protein entries in OpenProt compared to UniProtKB/TrEMBL, ionbot’s identification rate only decreased by ∼2% under FDP ∼1%. We show that different subgroups of peptides present different proportions of false identifications, which can be harmonised with entrapment-based filtering. However, if non-canonical identifications are not rigorously filtered, this can lead to many false matches, consistent with a small proportion of high-quality non-canonical PSMs in non-HLA proteomics [50,51]. This can be explained by a combination of short length, low abundance, and sample complexity.

Furthermore, the open search mode in ionbot drastically increased the identification rate compared to both FragPipe and ionbot run in closed search mode. This is consistent with previous observations that roughly one-third of all unidentified spectra in a classical LC-MS/MS setup originate from peptides carrying unexpected modifications [8].

Several PTMs observed on non-canonical peptides are unexpected. However, due to the low abundance of non-canonical proteins, we cannot determine if they are genuinely subject to extensive modifications, or if the high prevalence of modified peptides is simply a result of the advantages of open modification search in retrieving spectra. Nonetheless, having consistently detected modified versions of non-canonical proteins, we believe it is important to further characterize the PTMs of these proteins.

Our detailed comparison of closed and open search shows that 38% of matches to non-canonical peptides in closed search are better explained by canonical peptides with an unexpected modification in open search. This indicates that open search in combination with a proteogenomics database can actively correct misidentifications [52], as it constitutes the closest approximation possible to a complete search space. Thus, our results suggest that non-canonical proteins identified in closed search should undergo open search reanalysis to probe whether identification still holds. This capability extends beyond modifications and allows capturing protein variants even when they are absent in databases, by identifying mass shifts that mirror isobaric amino acid substitutions. Our analysis demonstrated that 3% of proteotypic PSMs’ mass shifts were consistent with alternative sequences absent from the initial search space. Combining proteogenomics with open search also allows for the consideration of mutated or modified versions of canonical peptides. The validity of peptide variants can be assessed by examining isobaric post-translational modifications (PTMs) after the search has been conducted [53].

A key limitation of ionbot is that it allows for only one unexpected modification per peptide. This constraint avoids the combinatorial explosion that comes from PSMs with unexplained mass shifts but reduces the tool’s ability to identify peptides with multiple unexpected modifications. In contrast, other open modification tools, such as MSFragger or SAGE, allow for more extensive modification matching, but at the cost of increased risk for overfitting and false identifications.

Moreover, even with careful search settings, many MS/MS spectra are still difficult to interpret with certainty. This is because different peptides can produce highly similar fragmentation patterns, leading to spectral ambiguity, a fundamental issue within mass spectrometry-based proteomics that is especially relevant in the context of proteogenomics open searches. Finally, modifications isobaric to SAAVs (e.g., methylation and Serine-to-Threonine) pose a challenge that cannot be solved through spectra analysis alone. In the future, retention time [55,56] and ion mobility [57,58] predictors could improve our ability to resolve these isobaric variations.

## Conclusions

Identifying non-canonical proteins necessitates not only the use of expanded sequence databases but also search strategies capable of addressing both sequence variants and post-translational modifications. Our findings indicate that combining proteogenomics with an open modification search is an effective strategy for discovering non-canonical peptides and significantly enhances the reproducibility of identifications.

At the same time, increasing the search space introduces new challenges in FDR control. We confirm a previous observation that when database search results include peptides with different properties (i.e., canonical and non-canonical), FDR levels should be assessed separately for each category. Our analysis of FDP produced contradicting results, as grouping led to lower FDP among non-canonical peptides, but higher FDP among canonical ones. Still, we encourage the community to investigate this aspect more closely.

Despite these challenges, our study affirms that the latest generation of open modification search engines can navigate the complexity of open modification proteogenomic search spaces and provides a practical and scalable path toward capturing the full complexity of the proteome.

## METHODOLOGY

### Datasets

**Table 3.**
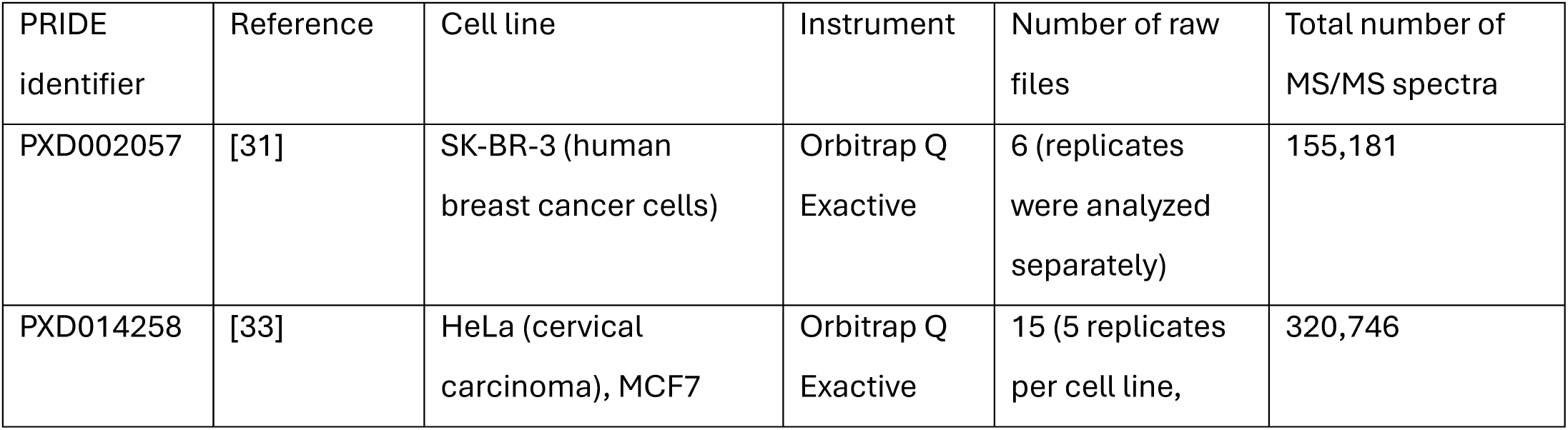
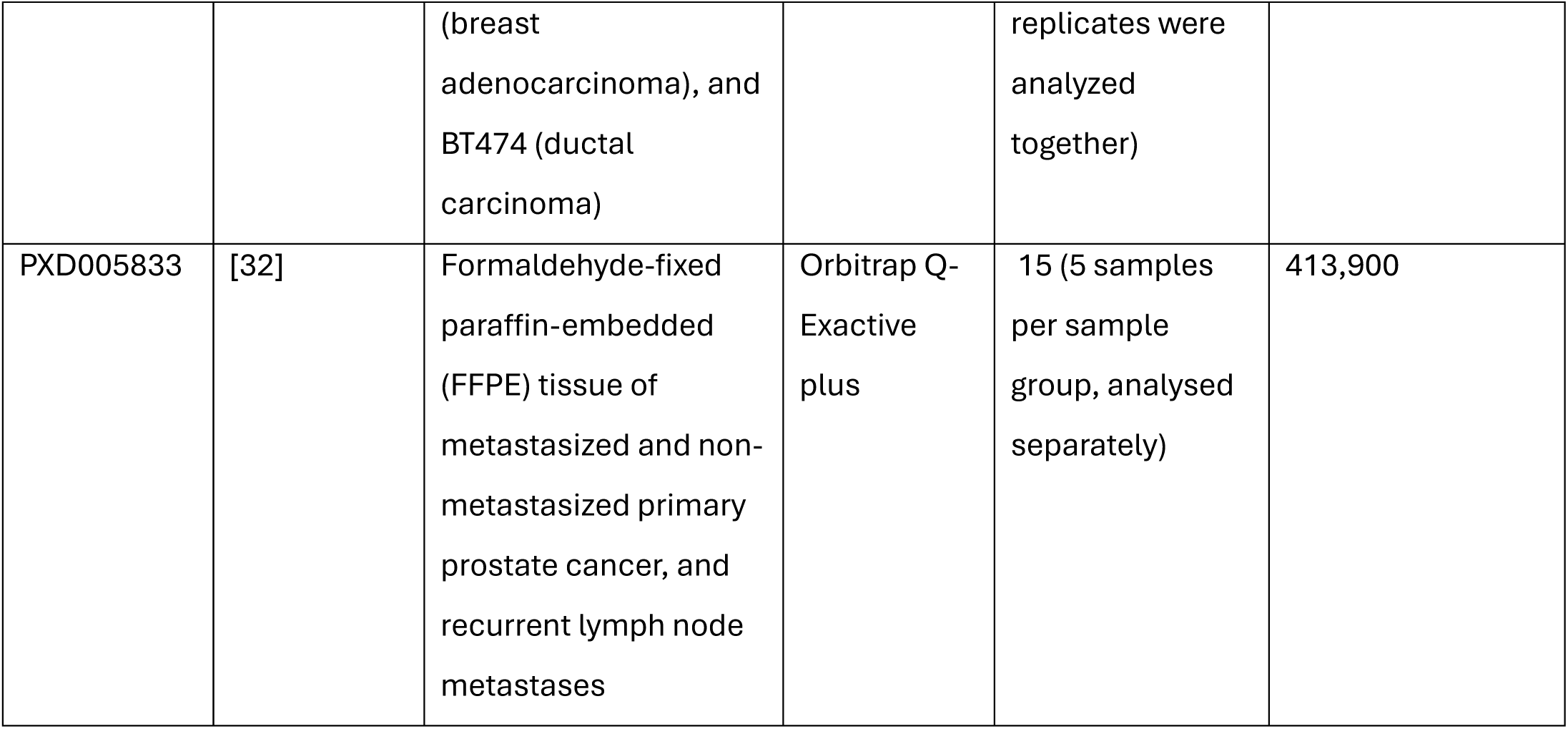
Public datasets.

| PRIDE identifier | Reference | Cell line | Instrument | Number of raw files | Total number of MS/MS spectra |
| --- | --- | --- | --- | --- | --- |
| PXD002057 | [31] | SK-BR-3 (human breast cancer cells) | Orbitrap Q Exactive | 6 (replicates were analyzed separately) | 155,181 |
| PXD014258 | [33] | HeLa (cervical carcinoma), MCF7 | Orbitrap Q Exactive | 15 (5 replicates per cell line, | 320,746 |
|  |  | (breast adenocarcinoma), and BT474 (ductal carcinoma) |  | replicates were analyzed together) |  |
| PXD005833 | [32] | Formaldehyde-fixed paraffin-embedded (FFPE) tissue of metastasized and non-metastasized primary prostate cancer, and recurrent lymph node metastases | Orbitrap Q-Exactive plus | 15 (5 samples per sample group, analysed separately) | 413,900 |

### Reference proteome databases

Raw data were searched against three protein databases: 1) Swiss-Prot reviewed reference proteins (referred to as the “Swiss-Prot” database, 20,419 entries); 2) Swiss-Prot reference proteins, their isoforms and TrEMBL unreviewed sequences (referred to as the “TrEMBL” database, 103,811 entries); 3) Swiss-Prot and TrEMBL sequences plus OpenProt non-canonical proteins (referred to as the “OpenProt” database, 683,413 entries). OpenProt proteins were considered non-canonical if their accessions started with II_ (novel isoform) or IP_ (alternative protein). UniProt version 2023_01 and OpenProt version 2 were used to construct the databases. Due to version differences, 2.6% of TrEMBL sequences do not overlap with Swiss-Prot and OpenProt, and 0.5% of Uniprot sequences do not overlap with OpenProt. Databases were merged with common contaminants from cRAP (https://www.thegpm.org/crap/). Decoys were generated with ionbot by reverting the target sequences, keeping Methionine at the starting position.

### Non-entrapment database search parameters and post-processing

For all database searches, the minimum and maximum peptide length were set to 7 and 30 amino acids, respectively. The peptide mass range was set from 500 to 6000 Da. The cleavage pattern was set to strict Trypsin, and up to two missed cleavages were allowed. N-terminal methionine clipping was enabled. The following modifications were indicated as variable carbamidomethylation of cysteine (+57.0215 Da), oxidation of methionine (+15.9949 Da), and N-terminal acetylation (+42.0106 Da). The mass tolerance was set to 20 ppm for both precursor and fragment ions.

Ionbot (version 11.4) was run in a Docker environment. In open PTM search mode, ionbot was allowed to match peptides with up to one of the PTMs listed in the UniMod database [36] (i.e., “unexpected” modifications) alongside any of the previously defined variable (expected) modifications.

Comet (TPP version 7.0.0) and MSFragger (version 4.2, FragPipe-23) search engines were run separately on samples from datasets listed in Table 3. For dataset PXD014258, the pepXML files were combined by cell line. PSMs were rescored with PeptideProphet [10], and proteins were inferred with pout2prot 1.2.1 (see below); PTM-Shepherd was used for mapping mass shifts to unimod annotation of PTMs for open search with FragPipe. More detailed information on parameters can be found in the repository (see Data availability).

FDR filtering was disabled during the search. Instead, q-values were recalculated on the output data using the “qvalues” function from the pyteomics Python package [54,59]. Identifications were ranked according to search engine scoring: by PSM score (descending) for ionbot, and by PEP (ascending) for TPP and FragPipe. PEP values were computed as 1−PeptideProphet probability. The formula used to calculate global FDR was:

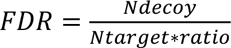

The value of the *ratio* parameter was set to 1.

Additionally, we performed group-wise FDR estimation. Briefly, we categorized the peptides into two subgroups based on whether the identified peptides corresponded to canonical or non-canonical proteins. A peptide was considered canonical if it mapped to at least one canonical protein. Conversely, a peptide was considered non-canonical if it mapped only to non-canonical proteins. Q-values specific to each subgroup were calculated using the pyteomics Python package (“group-wise” q-value) and the GroupWalk R package [22].

In non-entrapment runs, PSMs were filtered for global q-value < 0.01 and peptides for entrapment-based filter for each peptide type (canonical/non-canonical).

To determine the proteins identified from each search engine run, we used an in-house script based on pout2prot to generate a list of protein groups starting from the list of identified filtered peptides [60]. Highly repetitive proteins (Major Histocompatibility Complex components, antibodies) were excluded from this analysis to avoid unsustainable run times. Protein-level FDR was not calculated. Non-canonical protein groups were removed, and only proteins with at least one proteotypic peptide were considered.

### FDP assessment with entrapment search and entrapment-based filtering

The entrapment databases (for Swiss-Prot, TrEMBL, and OpenProt) were generated by shuffling the target peptide sequences using FDRBench (version 0.01) [30]. The entrapment ratio was set to 1. The databases were digested before shuffling. The digestion parameters were the same as for non-entrapment searches. Decoys were created by reversing both target and entrapment peptides, keeping Methionine at the starting position (if present). The three datasets, described above, were analysed with the resulting entrapment databases using ionbot (in both open and closed search mode), TPP, and FragPipe. The FDP (paired method) was calculated for peptide-level identifications from q-values, obtained through three different methods (global, group-wise, and group-walk), with FDRBench default parameters. Finally, peptide degeneracy was assessed and figures generated using in-house scripts available at https://github.com/Labo-MAB/Ionbot_FDP.

We concluded that an “entrapment-based” filtering worked best. In this approach, the maximal rank of peptide scores corresponding to FDP 1% for each peptide type was recorded for each sample. Next, peptide identifications in non-entrapment runs were sorted by PSM score and ranked, setting a threshold to the rank from entrapment runs.

### Analysis of modifications isobaric to amino acid substitutions

We selected peptides from the ionbot open search if at least one of their peptidoforms contained a modification isobaric with an amino acid substitution. As mass shifts often can be mapped to several PTMs, we only selected peptides containing the residues of expected substitutions. Considering that PTM localisation algorithms are imperfect [61], we assessed for variants of all residues on a peptide with expected substitutions (e.g., if a peptide with isoSAAV Alanine→Glutamine contains two Alanine residues, both are queried; if no Alanine residue is present, the peptide is ignored). Next, we mapped target residues (potential substitutions) to the genomic coordinates (GRCh38), where possible, and extracted their variant information from gnomAD (version 4.1.0) [39]. If the protein accession was absent in OpenProt 2.0 GTF, the genomic coordinates could not be mapped. If a codon, retrieved from the reference genome, did not encode a target residue, it was not queried in gnomAD. Target codons were retained if one base substitution (single-nucleotide variant) could lead to a target missense mutation.

### Non-canonical protein analysis according to HPP guidelines

We followed HPP guidelines version 3 (2019). We filtered out peptides derived from non-canonical proteins, identified with ionbot open search, if they were: 1) less than nine amino acids long, and 2) nested within another longer peptide. From the resulting set, we further filtered out partially overlapping peptides if their total length was less than eighteen amino acids. Retained peptides were checked for uniqueness using the HPP uniqueness checker web tool (https://hppportal.net/toolchecker.html). Peptides mapped to any tryptic peptide were filtered out. The resulting set of peptides was checked on potential SAAVs, if at least one of their PSMs contained a mass shift that was isobaric with amino acid substitution (as described above). Whenever the corresponding SAAVs were found in gnomAD, the spectra were matched to alternative peptides and these alternative PSMs were compared to the original PSMs with isoSAAVs. The Pearson correlation coefficient (PCC) of the assigned fragment ion intensities was compared to the intensities of predicted fragmentation spectra for the corresponding sequences of such pairs with MS2PIP v3.11.0 (HCD 2021 model) ion intensity prediction [63]. The mass shifts of the original peptides were preserved for alternatives, too. If multiple mass shifts were detected per peptide, the alternative peptidoform (with up to one mass shift masked) with the highest PCC was chosen per alternative peptide. The remaining PSM’s mass was checked to match another peptide in the database with one amino acid difference and a mass difference of less than 0.001 Da (none such cases were found). The protein was further considered if at least two proteotypic peptides were detected in any of the analysed datasets. To assess a protein’s ability to be detected according to HPP guidelines, we performed *in silico* digestion of the OpenProt v2 database and applied the same filtering criteria as described above. *In silico* digestion was done with Kiwi (https://github.com/Francis-B/Kiwi) with the same digestion parameters as for the abovementioned analyses.

### Visualization and examination of spectra identified by ionbot open search

Best-scoring PSMs of peptides that passed HPP guidelines were selected for visualisation and manual examination. Spectra were retrieved from mzXML files with the pyteomics Python package and visualised with spectrum_utils [64]. Theoretical intensities were predicted with MS2PIP, accounting for modifications. The PCC was calculated between the intensities of observed and predicted annotated spectrum peaks. When any peak was absent, its intensity was set to zero. Three researchers were involved in manual spectra examination, where two examined all selected spectra, and a third only those spectra with a controversial evaluation from the previous two. The list of examined PSMs, as well as their Universal Spectrum Identifiers (USI) and the scores from the examiners, can be found in Supplementary Table S2.

## Abreviations

LC-MS/MS: Liquid Chromatography - Tandem Mass Spectrometry
ORF: Open Reading Frame
OMS: Open Modification Search
PEP: Posterior Error Probability
PTM: Post-Translational Modification
PSM: Peptide-Spectrum Match
FDR: False Discovery Rate
FDP: False Discovery Proportion
TPP: Trans Proteomic Pipeline
SAAV: Single Amino Acid Variant
IsoSAAV: mass shift isobaric to SAAV
PCC: Pearson Correlation Coefficient
HPP: Human Proteome Project
USI: Universal Spectrum Identifier

## Ethics approval and consent to participate

Not applicable.

## Availability of data and materials

The datasets analysed during the current study are available in the PRIDE Archive repository [53] with dataset identifiers PXD002057, PXD005833, and PXD014258. The search engine output files as well as gnomAD uniqueness check results, are available on Zenodo (doi: 10.5281/zenodo.18023698). The Python code used to generate the results in this manuscript is available at the following link: https://github.com/CompOmics/oui-discovery

## Authors’ contributions

Study design: MAB, LM; Data generation: EM, VV, FB, SL; Data analysis and interpretation: EM, VV, TC, IA, MAB, LM; Writing and editing: VV, EM, TC, MAB; Funding: MAB, LM. All authors read and approved the final manuscript.

## Acknowledgements

We thank all members from the Martens and Brunet lab for helpful discussions. Computations were made in part on MAB’s HPC server maintained by the Centre de Calcul Scientifique de l’Université de Sherbrooke, and on the supercomputers Béluga and Narval, managed by Calcul Québec and the Digital Research Alliance of Canada. The operation of these supercomputers is funded by the Canada Foundation for Innovation (CFI), Ministère de l’Économie et de l’Innovation du Québec (MEI) and le Fonds de Recherche du Québec (FRQ).

